# A machine-learning-guided hydrogen-bonded organic framework for long-term, ultrasound-triggered pain therapy

**DOI:** 10.1101/2025.08.11.669645

**Authors:** Weilong He, Cathy Yang, Xi Shi, Yanxing Wang, Wenliang Wang, Alexander Schafer, Brinkley Artman, Liwen Zhou, Xiangping Liu, Kai Wing Kevin Tang, Jinmo Jeong, Zhecheng He, Henry Garcia, Alexa Olivarez, Erin H. Seeley, Sihan Yu, Anakaren Romero Lozano, Pengyu Ren, George D. Bittner, Huiliang Wang

**Author notes:** These authors contributed equally to this work (Weilong He, Cathy Yang, Xi Shi, Yanxing Wang).

## Abstract

Effective treatment of chronic pain remains hindered by the lack of drug delivery systems that simultaneously achieve long-term stability, high spatial precision, and non-invasiveness^1,2,3^. Here, we utilize a programmable, ultrasound-responsive drug delivery platform based on hydrogen-bonded organic framework (HOF) nanoparticles, enabling on-demand anesthetic release with long-term and durable analgesic efficacy. A machine learning (ML)–guided screening pipeline was developed to evaluate about 250 FDA-approved drugs, spanning both hydrophilic and lipophilic agents, and identified bupivacaine (lipophilic) and lidocaine hydrochloride (hydrophilic) as optimal candidates. These agents were efficiently encapsulated into HOF nanoparticles via diffusion and double-solvent methods. Ultrasound-triggered drug release *in vitro* transiently suppressed calcium signaling in adeno-associated virus (AAV)-transfected neurons. *In vivo*, ultrasound-activated release of bupivacaine or lidocaine HCl at the sciatic nerve site significantly elevated mechanical nociceptive thresholds for up to seven days, reduced autotomy (self-mutilaion) behavior, and improved motor function in a rat model of chronic pain. Notably, the machine learning–identified top candidates not only exhibited high loading efficiency, but also demonstrated superior therapeutic outcomes *in vivo*, establishing a direct link between computational prediction and biological efficacy. This non-invasive, programmable HOF-based system provides a clinically translatable platform for on-demand, spatiotemporally precise pain management.

## Introduction

Effective management of pain remains hindered in part due to the lack of drug delivery systems that combine high spatiotemporal precision, long-term efficacy, and non-invasiveness.^1,2^ In particularly, chronic pain affects over 20% of adults worldwide and 6.9% of adults have experienced high-impact chronic pain, leading to major healthcare costs and decreased productivity^4,5^. Among chronic pain conditions, peripheral nerve injury (PNI) affects hundreds of thousands annually^6–8^. Surgical repair often fails to restore full function, and systemic analgesics carry risks of addiction, adverse effects, and poor spatial specificity^9–11^. Patients frequently develop neuropathic pain and motor deficits, including self-mutilation and impaired locomotion^12–16^, underscoring the need for localized, on-demand drug delivery^17^.

While conventional analgesics—including non-steroidal anti-inflammatory drugs and opioid-based medications—provide rapid symptom relief, their long-term use raises serious concerns regarding addiction, tolerance, and systemic side effects^18–20^. In particular, the widespread use of opioids, though therapeutically effective, has contributed to a growing public health crisis due to misuse and dependency. The urgent need for safer, non-addictive alternatives has spurred growing interest in non-pharmacological and localized pain management approaches, such as electrical stimulation, cooling therapies, and optogenetic neuromodulation. However, these strategies remain limited in their clinical applicability due to invasiveness, complexity, or short duration of action. Recent insights into pain mechanisms and recovery pathways have emphasized the need for next-generation therapeutic modalities that integrate precise spatiotemporal control, extended drug stability, and minimal invasiveness to improve both efficacy and patient quality of life^21^.

To address these unmet needs, next-generation drug delivery platforms must not only ensure long-term drug stability and sustained release but also provide spatiotemporal control over therapeutic action^22,23^. In particular, the integration of neuromodulatory capabilities—such as externally triggered, localized drug release—enables precise inhibition or activation of neural circuits, offering significant advantages for treating pain and other neurological disorders^24,25^. Among various modalities, ultrasound provides a promising non-invasive trigger for deep-tissue drug release, allowing on-demand modulation with minimal systemic exposure. By combining prolonged retention, responsive release, and precise targeting, such systems can significantly improve patient autonomy and therapeutic outcomes in chronic disease management^24,26–29^.

Currently, most ultrasound-sensitive platforms rely on gas-filled microbubbles or thermosensitive lipids, which exhibit poor stability and are difficult to precisely control *in vivo*^30,31^. Recently, hydrogen-bonded organic frameworks (HOFs) have emerged as promising metal-free, biocompatible, and highly porous materials for drug delivery^32–35^, featuring dynamic π–π stacking and hydrogen bonding interactions that enable high drug-loading capacity (∼30 wt%) and mechanical responsiveness^36–40^. However, three critical challenges remain for advancing HOFs toward clinically meaningful pain therapeutics. First, the encapsulation of hydrophobic drugs in HOFs has not been demonstrated for potent anesthetics or neuromodulators. Many clinically important pain therapeutics are highly lipophilic—facilitating membrane penetration but also causing poor aqueous solubility, rapid systemic clearance, and off-target effects.^41^ Achieving high-capacity loading together with long-term retention and spatiotemporally precise release would greatly expand the therapeutic scope of HOFs. Second, no systematic, predictive framework exists to identify drugs most compatible with a given HOFs structure for specific biomedical applications. Current studies rely on trial-and-error screening, which is time-consuming, resource-intensive, and prone to overlooking high-potential candidates. Third, ultrasound-activated drug release for pain management has not been validated in clinically relevant injury models. Most studies are conducted in healthy animals, which fail to replicate the inflammatory, neuropathic, and motor-deficit environment following nerve injury^41,42^.

To address key limitations in current pain management platforms, we developed a HOF-based nanocarrier system featuring high drug loading capacity, structural stability, and precise ultrasound-triggered release. Specifically, we: (**i**) demonstrate, for the first time, the encapsulation of a hydrophobic anesthetic within HOF–TATB; (**ii**) establish a machine learning (ML)–guided screening pipeline to identify drugs with optimal loading and release characteristics; and (**iii**) validate ultrasound-activated, on-demand anesthetic release in a rodent model of neuropathic pain. The solid-state, modular design of HOF–TATB enables tunable drug encapsulation while preventing premature leakage and maintaining ultrasound responsiveness. Our ML-driven approach evaluates drug physicochemical properties to predict compatibility, while versatile loading strategies accommodate a broad range of drug chemistries. By uniting AI-guided drug selection, high payloads, and spatiotemporally precise release, this platform offers a non-invasive, controllable solution for long-term pain relief and holds broader potential for neuromodulation and neurological disorder therapies (Fig. 1a).

**Fig. 1.**
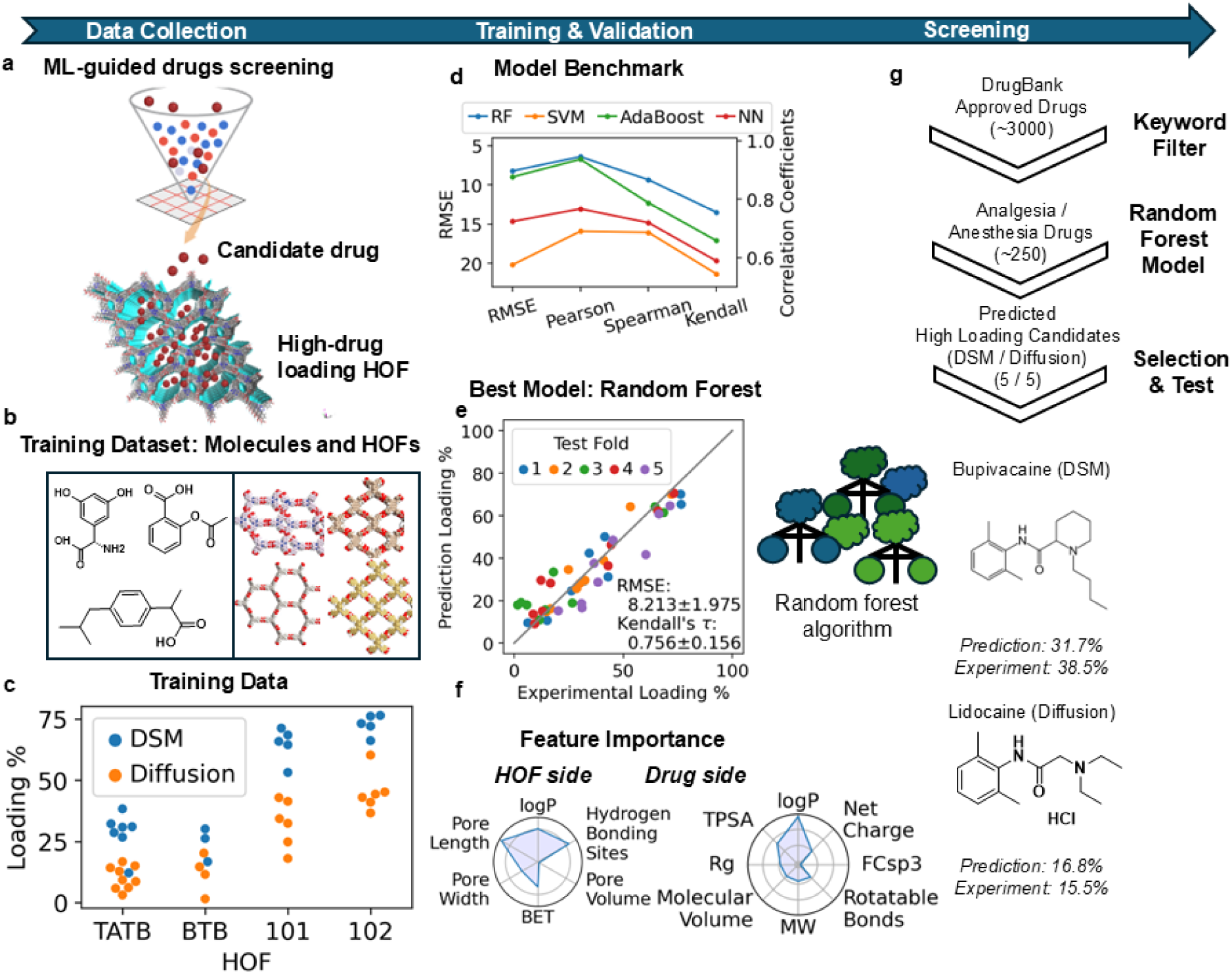
Schematic illustration of the machine learning–guided design and FUS triggered local anesthetic (LA) delivery platform for precision pain management. (a) Machine learning–guided screening of drug molecules for high-loading hydrogen-bonded organic frameworks (HOFs). Candidate drugs predicted to exhibit high loading are selected for experimental validation with representative HOF materials. (b) Left: Representative small-molecule drugs used for model training and screening. Molecular descriptors included molecular weight, logP, TPSA, rotatable bonds, and net charge. Right: Structures of four representative HOF materials (HOF-TATB, HOF-101, HOF-BTB, and HOF-102) used in drug loading studies, with structural descriptors such as pore size, BET surface area, and number of hydrogen-bonding sites extracted as model inputs. (c) Experimentally measured drug loading capacities for 15 molecules across four HOFs using two loading strategies: double solvent method (DSM) and diffusion method. Each point represents a drug–HOF combination. (d) Benchmarking performance of four machine learning models—Random Forest (RF), Support Vector Machine (SVM), AdaBoost, and Neural Network (NN)—using RMSE and correlation coefficients (Pearson, Spearman, Kendall) under five-fold cross-validation. (e) Scatter plot comparing predicted vs. experimentally measured loading capacities using the best-performing model (Random Forest). The model achieves a mean RMSE of 8.213 ± 1.975 and a Kendall’s τ of 0.756 ± 0.156. (f) Feature importance analysis of the Random Forest model. On the HOF side, pore size, lipophilicity (logP), and hydrogen bonding were key predictors. On the drug side, lipophilicity (logP), total polar surface area (TPSA), and molecular volume were most influential. (g) Workflow of virtual screening pipeline. Approximately 3,000 FDA-approved drugs were filtered to ∼250 analgesics/anesthetics. The Random Forest model was used to predict loading capacity, followed by experimental validation of the top 5 candidates. Bupivacaine and Lidocaine HCl were ultimately selected based on model score, interaction energy, and translational potential.

## Results

### 1. Machine Learning–Guided Identification of High-Loading Anesthetic Candidates

Despite recent advances in nanocarrier-based drug delivery, the identification of optimal therapeutics for a given delivery platform remains largely empirical. In the context of HOFs, drug loading efficiency is governed by a non-trivial interplay of molecular properties, including lipophilicity (logP), hydrogen bonding potential, molecular size, and steric compatibility. Conventional screening relies on labor-intensive, trial-and-error experimentation, which limits scalability and slows translational progress.

Recognizing the limitations of empirical screening, we sought to establish a predictive model using quantitative descriptors that reflect the underlying drug–HOF interaction mechanisms. We carefully crafted a few structural descriptors for HOFs and drugs as the input of the models based on our preliminary understanding that non-specific interactions such as hydrophilicity was the main force during the process of loading drug into HOF crystals. The descriptors we used for drugs were the volume of molecule, the radius of gyration, molecular weight, the number of rotatable bonds, the ratio of sp^3^ hybridization carbons, total polar surface area, logP, and the net charge at pH 7 (Fig. 1b left). The descriptors we used for HOFs were the number of hydrogen bonding sites, crystal pore volume, crystal port size, crystal BET surface area, and logP (Fig. 1b right). All non-crystal features were computed with RDKit^43^.

To identify analgesic and anesthetic drugs with high loading capacities for HOFs, we curated a focused library of candidate molecules. From the ∼3,000 FDA-approved molecules in the DrugBank^44^ database, we extracted a subset of ∼250 compounds annotated for analgesic or anesthetic activity. (SI Tab.1 Focused Library) This collection was screened using our machine learning–based virtual pipeline, enabling a rapid and cost-effective alternative to conventional experimental testing.

Our training dataset was curated by experimentally measuring the drug loading capacities across four distinct HOF materials for 15 chemically diverse small molecules, including commonly available drugs and dyes. This yielded a total of 45 drug–HOF combinations as training data (Fig. 1c, SI Tab.2 Training Dataset). Using this dataset, we trained and benchmarked several machine learning algorithms—Support Vector Machine (SVM)^45^, Random Forest (RF)^46^, AdaBoost^47^, and Neural Network (NN)^48^—under a 5-fold cross-validation scheme (Fig. 1d). Among these, the Random Forest model achieved the best predictive performance, exhibiting the highest correlation coefficients and the lowest root-mean-square error (RMSE) across all test folds (Fig. 1e). The corresponding scatter plot (Fig. 1e) illustrates strong agreement between predicted and experimental values, with an overall RMSE of 8.213 ± 1.975 and a Kendall’s τ of 0.756 ± 0.156. We placed particular emphasis on Kendall’s τ, as it quantifies the model’s ranking ability—a key requirement for identifying top-performing drug candidates based on relative loading potential. To interpret model decisions and elucidate feature contributions, we performed a permutation-based feature importance analysis^49^ (Fig. 1f). On the HOF side, pore size and lipophilicity (logP) emerged as the most influential descriptors. On the drug side, logP and total polar surface area (TPSA) had the strongest impact on prediction accuracy. These results suggest that the model effectively captured the dominant roles of spatial compatibility and hydrophobic interactions in governing drug loading into HOFs.

Given its superior performance, the Random Forest model was selected to screen the focused drug library (Fig. 1g). To enhance prediction robustness and generalizability, we employed an ensemble strategy, averaging the outputs from the five models generated during cross-validation to derive the final loading score for each compound. Following this screening, the top 5 candidates for HOF-TATB were shortlisted for further evaluation using either encapsulation strategy. Final selection was guided by both theoretical predictions and practical considerations, including commercial availability, cost, established mechanism of action, and predicted drug–HOF interaction energy—a surrogate metric for formulation stability (SI Fig.1 Top 5 Candidates).

Among these potential drug candidates, bupivacaine and lidocaine HCl emerged as the most promisin. Bupivacaine was prioritized due to its consistently strong predicted interaction with the HOF-TATB framework and its long-acting anesthetic profile, which aligns with our goal of sustained local pain relief. Although amylocaine showed slightly higher predicted interaction, lidocaine HCl—a widely used local anesthetic with rapid onset but short half-life (∼90 min), hydrophilic nature, and rapid systemic clearance—was ultimately selected based on its excellent clinical safety, rapid onset of action, lower cost, and wide commercial availability, which are key factors for translational feasibility. These properties, however, also limit their utility for sustained or programmable analgesia, often requiring frequent re-administration and posing a risk of systemic toxicity. These ML predictions guided our subsequent experimental formulation and validation efforts.

#### Synthesis and Characterization of HOF@Lidocaine HCl & Bupivacaine

To experimentally validate the machine learning–guided selection of high-loading anesthetic candidates, we synthesized HOF-TATB and encapsulated the top two predicted drugs—lidocaine hydrochloride (hydrophilic) and bupivacaine (lipophilic)—into its porous framework. Among several candidate materials, HOF-TATB, assembled from 4,4′,4″-s-triazine-2,4,6-triyl-tribenzoic acid (H TATB), was selected for its structural robustness, colloidal stability (SI Fig. 2), and strong responsiveness to ultrasound stimulation^50^ (Fig. 1b right). The framework is stabilized through directional hydrogen bonding and π–π stacking interactions, which facilitate efficient drug encapsulation and enable mechanical energy–triggered release.

To accommodate differences in drug hydrophilicity, distinct loading strategies were adopted (Fig. 2a): lidocaine HCl was loaded via aqueous diffusion, leveraging multivalent hydrogen bonding and π–π stacking interactions to achieve efficient encapsulation (∼9.8 wt%) and stable retention under physiological conditions. In contrast, bupivacaine was incorporated using a dual-solvent approach, involving interfacial diffusion from a dichloromethane (DCM) solution into a water-dispersed HOF suspension, achieving a high loading efficiency of approximately 30.0 wt%. Transmission electron microscopy (TEM) revealed well-defined, crystalline nanoparticles for both formulations, with diameters ranging from ∼400 to 500 nm (Fig. 2b), confirming the preservation of structural integrity post-loading. Powder X-ray diffraction (PXRD) further verified that the crystallinity of HOF-TATB was retained after drug encapsulation (Fig. 2c), indicating that guest incorporation does not compromise framework order. Experimental quantification of drug loading revealed that lidocaine HCl and bupivacaine achieved high loading efficiencies (SI Fig. 3 and SI Fig. 4), in close agreement with machine learning predictions (Fig. 2d). This consistency underscores the practical utility of integrating AI-guided selection with modular HOF nanocarriers to streamline formulation design for ultrasound-triggered drug delivery. Collectively, these results demonstrate that ML-prioritized anesthetic candidates can be efficiently and stably encapsulated within structurally robust HOF systems, establishing a foundation for subsequent ultrasound-responsive release and *in vivo* therapeutic testing.

**Fig. 2.**
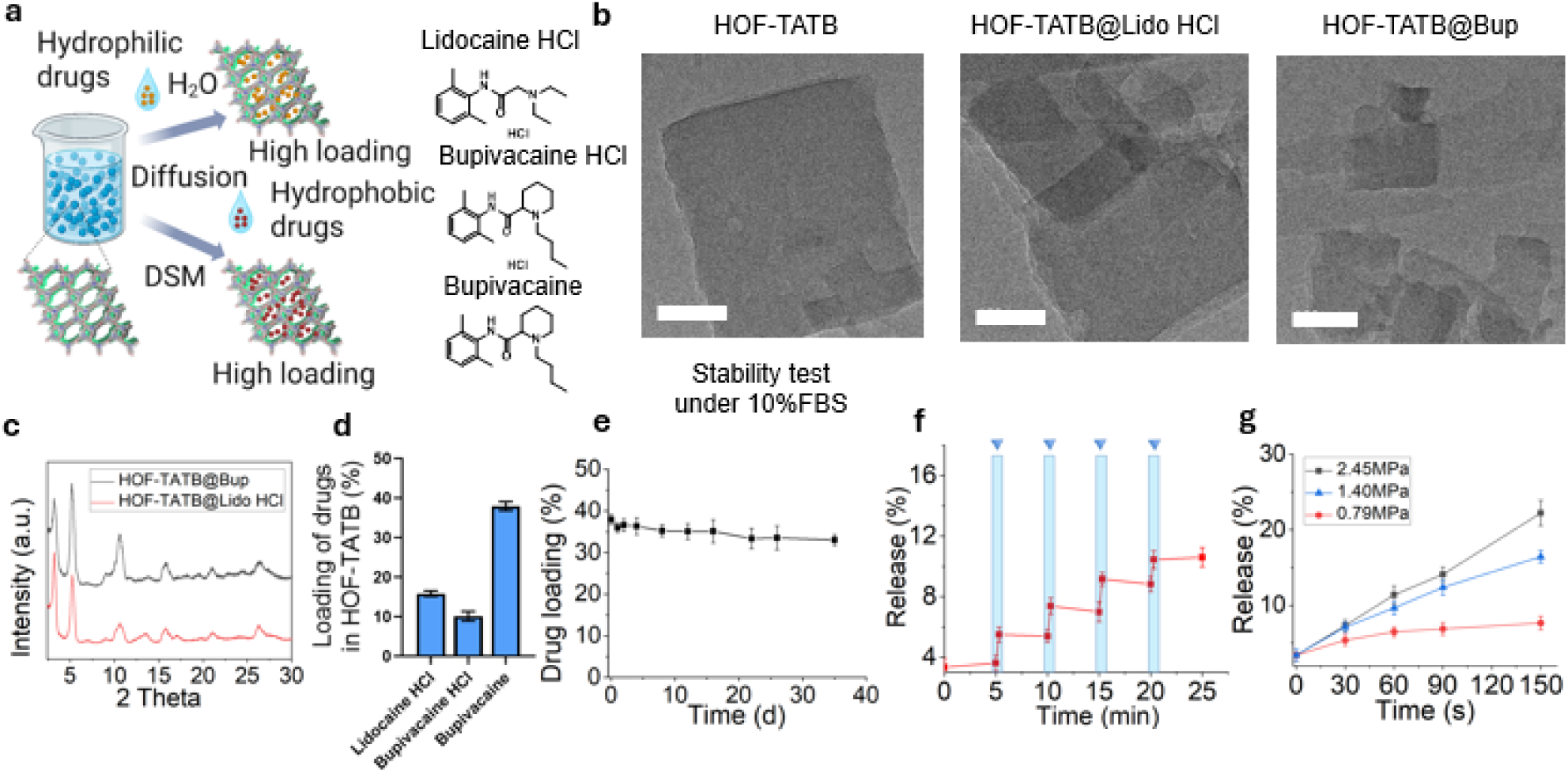
Stability and ultrasound-responsive drug release of HOF-TATB nanoparticles loaded with various therapeutics via the dual-solvent method. (a) Schematic illustration of two loading strategies for hydrophilic and hydrophobic anesthetic drugs into HOF-TATB— aqueous diffusion for hydrophilic drugs and a dual-solvent method (DSM) for hydrophobic drugs—enabling high loading capacity across diverse drug chemistries. (b) TEM images of HOF-TATB nanoparticles before and after loading with lidocaine HCl or bupivacaine, showing that the particle morphology and crystalline structure were retained after drug loading. (c) Powder X-ray Diffraction (PXRD) patterns of HOF-TATB nanoparticles before and after loading with bupivacaine or lidocaine HCl, showing that the characteristic diffraction peaks were preserved, indicating retention of the crystalline structure after drug loading. (d) Drug loading capacity of HOF-TATB nanoparticles for lidocaine HCl, bupivacaine HCl, and bupivacaine, showing the highest loading for bupivacaine due to its higher hydrophobicity. The experimental loading trends were consistent with the machine learning–predicted rankings. (e) Long-term storage stability of HOF-TATB@bupivacaine prepared via the dual-solvent method (DSM), showing minimal decline in drug loading over 30 days in 10% fetal bovine serum (FBS) at 25 °C. (f) Ultrasound-triggered release of bupivacaine from HOF-TATB nanoparticles *in vitro* under varying ultrasound intensities at 1.5 MHz, demonstrating a dose-dependent increase in drug release with increasing acoustic power (0.79MPa, 1.40MPa, 2.45MPa). All data are presented as mean ± s.d., n = 3 independent replicates. (g) Pulsatile release profile of HOF-TATB@Bup under repeated FUS exposures (1.5 MHz, 1.40MPa, 1 min pulses at 5-minute intervals), showing stepwise increases in cumulative release with each stimulus, indicating the feasibility of on-demand release.

#### *In vitro* Ultrasound-Triggered Drug Release and Stability Evaluation

Building on the machine learning–guided identification and successful encapsulation of lidocaine HCl and bupivacaine into HOF-TATB, we next investigated the stability and ultrasound-responsiveness of these drug-loaded HOFs to evaluate their potential for sustained and controllable anesthetic delivery. Drug retention under physiological conditions was first assessed by incubating HOF-TATB loaded with bupivacaine in 10% fetal bovine serum (FBS) at 25 °C. HOF-TATB loaded with bupivacaine showed little change in drug loading over 30 days in 10% FBS at 25 °C (Fig. 2e), indicating excellent stability in a protein-rich environment.

We then tested the ultrasound-triggered release profile of bupivacaine-loaded HOFs under varying acoustic power. The ultrasound-triggered release experiments demonstrated that the percentage of drug released increased with the ultrasound peak pressure applied (Fig. 2f), confirming the system’s capacity for on-demand, repeatable, and programmable dosing. Upon exposure to focused ultrasound, bupivacaine release increased with acoustic intensity, achieving 22.3 ± 1.7% release within 150 s under 2.45 MPa (Fig. 2f). In contrast, only 16.4 ± 0.8% release was observed under 1.40 MPa, highlighting the tunability of drug release with ultrasound strength. Moreover, bupivacaine release from HOF-TATB could be repeatedly triggered by sequential ultrasound pulses (20 s per pulse), demonstrating stepwise and cumulative drug release. Four successive activations led to incremental release levels of 5.5 ± 0.5%, 7.4 ± 0.5%, 9.2 ± 0.4%, and 10.4 ± 0.6%, respectively (Fig. 2g), highlighting the programmable and re-triggerable nature of this delivery system. In contrast to conventional passive release formulations, this ultrasound-triggered strategy enables dynamic dosing control in response to clinical needs.

#### *In vitro* Ultrasound-Triggered Drug Release and Neuron Evaluation

To validate the neuromodulatory functionality of our HOF-based nanocarriers, next we performed calcium imaging in cultured neurons. Such *in vitro* testing provides a critical proof-of-concept for evaluating the platform’s ability to modulate neuronal excitability in a spatially and temporally controlled manner, which is essential for future neurotherapeutic translation. We transduced primary neurons with AAV carrying GCaMP6s, enabling real-time optical recording of intracellular Ca² fluctuations as a proxy for neuronal activity (Fig. 3a and 3b).

**Fig. 3.**
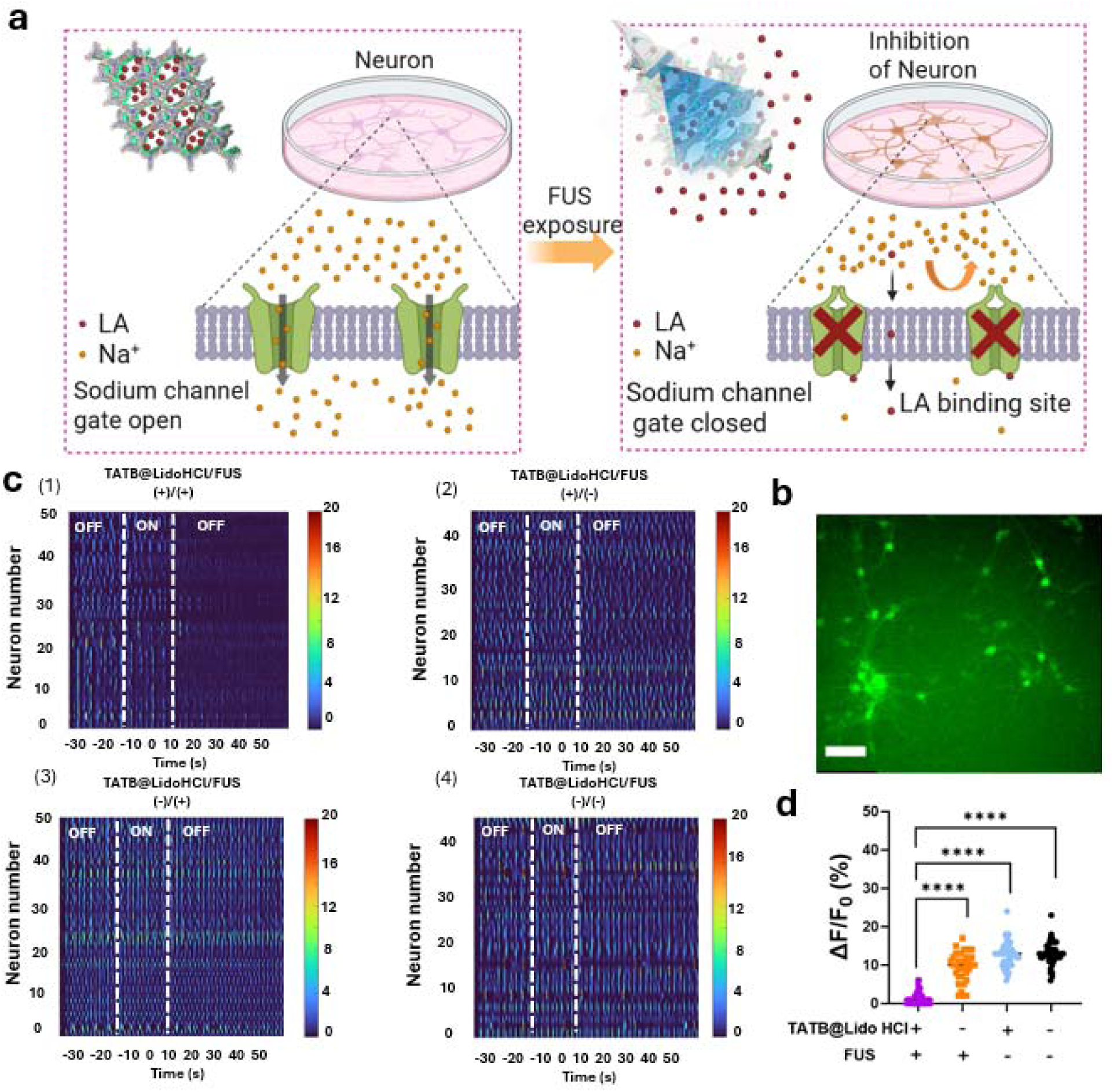
Ultrasound-triggered release of lidocaine from HOF-TATB nanoparticles and resulting inhibition of neuronal activity *in vitro*. (a) Schematic illustration of the mechanism by which FUS stimulation induces the release of lidocaine (LA) from HOF-TATB nanoparticles, enabling its interaction with voltage-gated sodium channels and leading to channel blockade. (b) Representative fluorescence image of primary cortical neurons expressing the calcium indicator GCaMP6s (hSyn::GCaMP6s-WPRE-SV40), used to monitor real-time neuronal activity. Scale bar, 50 μm. (c) Heat maps of normalized GCaMP6s fluorescence intensity from individual neurons under different experimental conditions (n = 40–50 neurons per group, three independent experiments), including (1) TATB@Lido HCl/FUS/(+)/(+), (2) TATB@Lido HCl/FUS/(+)/(-), (3) TATB@Lido HCl/FUS/(-)/(+) and (4) TATB@Lido HCl/FUS/(-)/(-). FUS (-) = ultrasound off; FUS (+) = ultrasound on (1.5 MHz, 1.40 MPa, 10 s per pulse). (d) Quantification of ΔF/F responses across neuron populations under control (TATB@Lidocaine, no FUS) and FUS-treated conditions. (****P < 0.0001, two-tailed unpaired t-test). All data are presented as mean ± s.d..

Neurons were first incubated with HOF-TATB@Lido HCl nanoparticles, during which robust and spontaneous calcium oscillations persisted, indicating that the encapsulated drug remained inactive under basal conditions (Fig. 3c). Upon exposure to a focused ultrasound pulse (1.5 MHz, 1.40 MPa, 10 s), we observed a rapid and widespread suppression of calcium transients across the majority of neurons at the experimental groups TATB@Lido HCl/FUS/(+)/(+) but not in other 3control groups (Fig. 3c–d). This outcome confirms effective ultrasound-triggered release of lidocaine from the HOF matrix, resulting in acute silencing of neuronal activity.

#### *In Vivo* Ultrasound-Triggered Analgesia and Programmable Pain Modulation

Building on these *in vitro* findings, we next evaluated the *in vivo* analgesic performance of HOF-TATB@lidocaine HCl. To this end, Sprague–Dawley rats were employed to establish a sciatic nerve block model, with ultrasound imaging–guided perineural injection of HOF-TATB@lidocaine HCl (Fig. 4a, SI Fig. 5 and SI Fig. 6). Following nociceptive threshold baseline acquisition, we systematically investigated the influence of FUS intensity on drug release from HOF-TATB@lidocaine HCl nanoparticles *in vivo* (1.5 MHz, 0.2 s on / 0.8 s off pulse cycles, total 8 min) (Fig. 4b). Strikingly, a relatively low intensity of 0.79 MPa effectively elicited a substantial and statistically significant increase in von Frey (VF) thresholds within 0.5 h post-FUS, indicating good analgesic effect. Higher intensities of 1.40 MPa and 2.45 MPa produced even more pronounced effects (p < 0.05), with analgesic efficacy persisting for up to 5h. The temporal profile of VF responses (Fig. 4c) further confirmed the dose-dependent and sustained analgesic effects of ultrasound, with peak responses observed within the first hour and a gradual decline over the next few hours. Notably, 1.40 MPa is well below the U.S. FDA limit for clinical diagnostic ultrasound intensity (I_SPTA_), highlighting its translational safety margin.

**Fig. 4.**
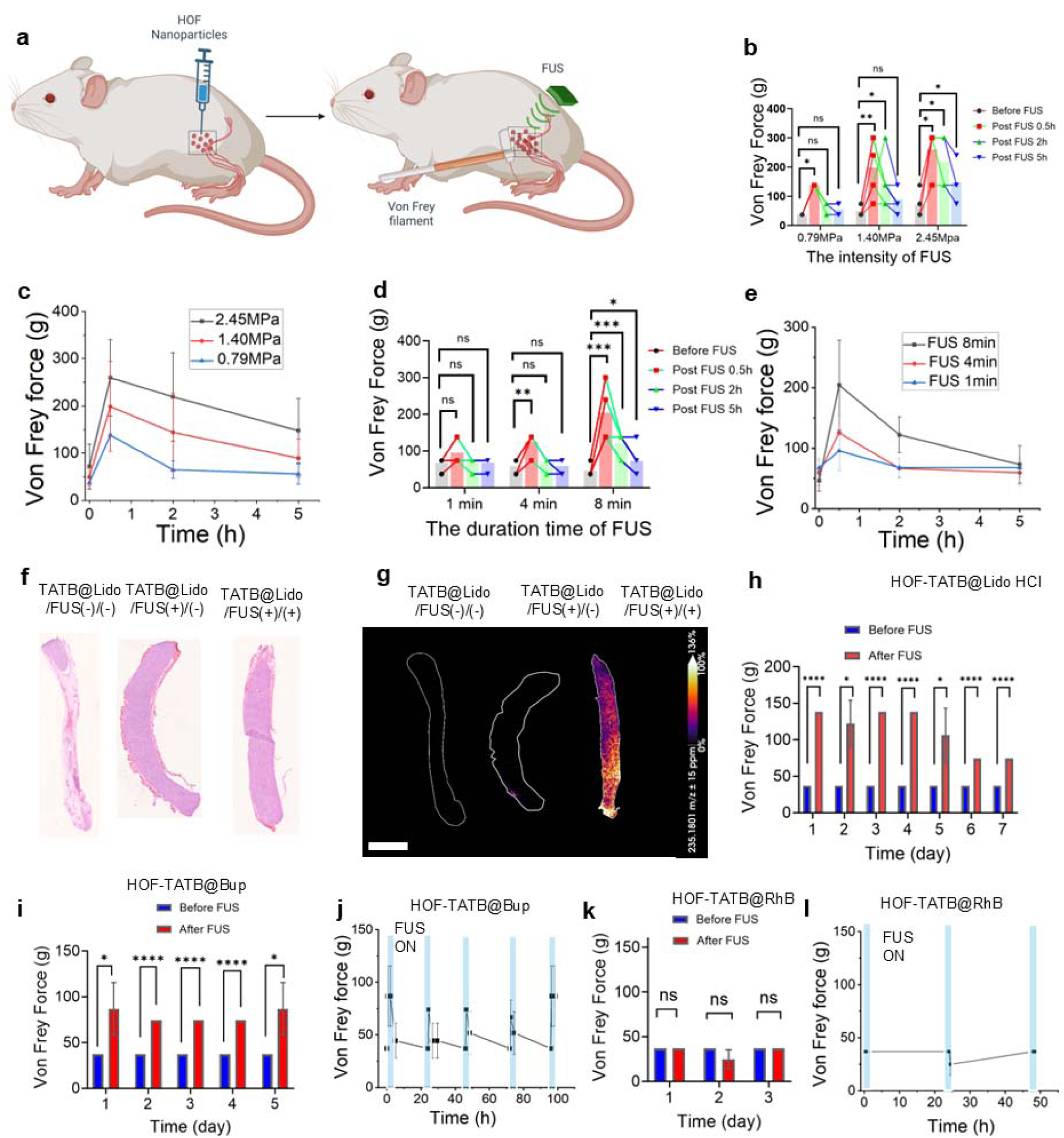
Ultrasound-triggered analgesia *in vivo* using HOF-TATB nanoparticles loaded with local anesthetics. (a) Schematic illustration of ultrasound-triggered analgesic delivery. HOF-based nanoparticles loaded with lidocaine HCl (TATB@Lidocaine HCl) were locally injected near the sciatic nerve of rats. Upon external FUS stimulation, the nanoparticles release lidocaine HCl to block nerve conduction, and mechanical nociceptive sensitivity was evaluated using von Frey filament testing. (b) von Frey forces before and after FUS at 0.79 MPa (n = 4), 1.40 MPa (n = 6), and 2.45 MPa (n = 4). Withdrawal thresholds increased markedly at 1.40 MPa and 2.45 MPa 0.5 h post-FUS (*p* < 0.05 vs. before FUS), with effects gradually declining over 5 h. Minimal changes were observed at 0.79 MPa. (c) Time-dependent changes in von Frey force after FUS. 1.40 MPa and 2.45 MPa groups showed a rapid increase in threshold at 0.5 h, followed by gradual decline. 2.45 MPa maintained the highest force throughout. (d) von Frey force before and after FUS with durations of 1 min (n = 6), 4 min (n = 5), and 8 min (n = 8). Significant increases in withdrawal thresholds were observed at 8 min 0.5 h post-FUS (*p* < 0.05– 0.0001 vs. before FUS), while minimal or no changes were detected at 1 min and 4 min. (e) Time-course of von Frey forces post-FUS at durations of 1 min (n = 6), 4 min (n = 5), and 8 min (n = 8). Only the 8-min FUS group showed a marked elevation in withdrawal threshold at 0.5 h, followed by a gradual decline over 5 h. No significant change was observed in the 1-min and 4-min groups. (f) H&E-stained sciatic nerve sections and corresponding MSI maps were obtained from three treatment groups (g): Sample 1 (no lidocaine HCl-loaded nanoparticles, no FUS), Sample 2 (with nanoparticles but no FUS), and Sample 3 (with nanoparticles and FUS). Lidocaine HCl accumulation was observed only in Sample 3 following FUS (1.5 MHz, 1.40 MPa, 0.2 s on / 0.8 s off, total 8 min). MSI signals at m/z = 235.1801 (±15 ppm) were normalized to total ion current. Scale bar: 3 mm. (h) Day-by-day comparison of nociceptive sensitivity thresholds before and after 8-min FUS in rats treated with HOF-TATB@lidocaine HCl. A significant increase in threshold was observed across all 7 days (n = 4, *P* < 0.05 to \*\*\**P* < 0.0001, paired t-test). (i) Statistical comparison of thresholds before and after FUS across 5 consecutive days for the HOF-TATB@bupivacaine group, confirming consistent and reversible analgesia (*P < 0.05 to **P < 0.01). (j) von Frey force changes in the HOF-TATB@Bupivacaine group following repeated FUS stimulation. Blue shaded areas indicate FUS application periods, which consistently increase mechanical thresholds at each stimulation time point, followed by gradual decline until the next FUS session. (k) Rats injected with Rhodamine b loaded HOF-TATB nanoparticles and stimulated with FUS served as nanoparticle-only controls, showing no elevation in mechanical thresholds (n = 3 rats). Corresponding statistical analysis for the HOF-TATB control group revealed no significant nociceptive changes (ns, not significant). (l) von Frey force in the HOF-TATB@RhB group. Blue shaded areas indicate FUS application; no significant threshold changes were observed. All data represent mean ± s.d. from biological replicates as indicated.

We therefore selected 1.40 MPa to examine the effect of FUS duration, revealing a clear time-dependent enhancement in nociceptive threshold from 1 min to 8 min stimulation (Fig. 4d-e). These findings demonstrate that both FUS intensity and duration can be tuned to achieve graded, on-demand anesthesia, offering a level of temporal precision currently unattainable with conventional local anesthetic injections. Clinically, such a noninvasive, remotely controllable system could enable personalized pain management—delivering brief, targeted analgesia for short procedures or sustained blockade for extended interventions—while minimizing systemic drug exposure and associated side effects.

To confirm that the observed behavioral analgesia resulted from localized, ultrasound-triggered drug release at the target site, we performed matrix-assisted laser desorption/ionization mass spectrometry imaging (MALDI-MSI) on longitudinal sciatic nerve sections (Fig. 4f-g and SI Fig. 7). MALDI-MSI confirmed that this behavioral improvement correlated with a robust and spatially confined accumulation of lidocaine at the targeted sciatic nerve site, observed only in the FUS-activated nanoparticle group (Fig. 4g).

Importantly, histological evaluation revealed preserved nerve morphology, underscoring the safety of this localized activation. Hematoxylin and eosin (H&E) staining of sciatic nerve sections 7 days post-treatment showed intact myelin structure and normal fiber morphology in both FUS-treated and untreated groups, with no evidence of degeneration, inflammatory infiltration, or tissue damage (SI Fig. 8). Immunofluorescence analysis further confirmed minimal Iba1-positive macrophage/microglia infiltration (SI Fig. 9) and negligible cleaved Caspase-3 staining (SI Fig. 10), indicating no significant immune or apoptotic responses. These results collectively support that TATB@Lidocaine HCl, with or without FUS activation, does not induce noticeable histopathological changes in sciatic nerve tissue.

To assess the reproducibility and stability of the ultrasound-triggered anesthetic effect, we administered daily FUS stimulation (1.40 MPa, 8 min) for 7 consecutive days following perineural injection of HOF-TATB@lidocaine HCl. VF test scores in Sprague Dawley rats consistently revealed a robust and statistically significant elevation in nociceptive threshold after each FUS session compared to baseline, with no apparent decline in efficacy across the treatment period (Fig. 4h). This indicates that the nanoparticle platform maintains its responsiveness to repeated ultrasound activation over multiple days, without evident tachyphylaxis or functional degradation. Such sustained, repeatable on-demand analgesia could be particularly advantageous in clinical scenarios requiring periodic pain relief—such as post-operative recovery or chronic pain flare-ups—while avoiding the need for continuous systemic drug administration.

Having established the feasibility of ultrasound-triggered release of hydrophilic anesthetics using lidocaine HCl, we next evaluated the long-term stability and therapeutic durability of our second machine learning–guided drug platform with a lipophilic anesthetic—bupivacaine. Due to its higher hydrophobicity and greater loading capacity (∼30.0 wt%), we hypothesized that bupivacaine would exhibit stronger retention within the hydrophobic HOF-TATB pore channels than lidocaine HCl, thereby enabling sustained *in vivo* localization and prolonged release upon ultrasound activation. To test this, Sprague–Dawley rats received ultrasound imaging–guided perineural injections of HOF-TATB@bupivacaine nanoparticles near the sciatic nerve. Using previously optimized FUS parameters (1.5 MHz, 1.40 MPa, total 8 min), we stimulated the nanoparticle depot once daily for five consecutive days and performed repeated von Frey tests following each activation. Animals injected with HOF-TATB@bupivacaine exhibited a robust and repeatable elevation in nociceptive threshold in VF tests after each FUS pulse, sustained across the entire five-day testing period (Fig. 4i–j). Compared with lidocaine HCl, bupivacaine produced similarly strong immediate analgesia and suggested more consistent multi-day repeatability, potentially attributable to its higher hydrophobicity, greater loading capacity, and stronger retention within the HOF-TATB framework. These results confirm the capability of our system for durable drug retention and reliable, on-demand analgesia over extended timescales. In contrast, control animals receiving unloaded HOF-TATB nanoparticles showed no significant change in mechanical sensitivity despite identical FUS stimulation (Fig. 4k–l), ruling out material-related or sonomechanical effects. Importantly, the consistent efficacy across multiple days underscores the long-term structural stability of the HOF nanocarrier *in vivo* and its potential for chronic pain applications requiring repeated intervention.

#### Modeling Neuropathic Pain and Motor Dysfunction after Sciatic Nerve Injury

To further assess the long-term therapeutic efficacy and clinical relevance of our ultrasound-responsive HOF nanoplatform, we employed an often-used rat model of sciatic nerve transection and repair—a well-established protocol that involves key hallmarks of chronic neuropathic pain, including spontaneous pain behavior and motor dysfunction.^51^ Following surgical transection and microsuture-based repair of the sciatic nerve, HOF-TATB nanoparticles encapsulating either bupivacaine or lidocaine HCl were perineurally injected at the injury site (Fig. 5a) once weekly, from week 2 to week 5. FUS (1.5 MHz, 8 min) was applied daily throughout this period to trigger on-demand anesthetic release. Motor function recovery was assessed using the Sciatic Functional Index (SFI), a widely adopted gait analysis metric that quantitatively primarily evaluates locomotor deficits following sciatic nerve injury (Fig. 5b).^51^ SFI values enable precise tracking of functional restoration over time. In parallel, toe-chewing (TC), self-mutilation behavior—a hallmark of spontaneous neuropathic pain in Sprague-Dawley rat—was evaluated as a robust indicator of dysesthesia severity (Fig. 5c).^15^ Compared to untreated controls, both HOF-TATB@bupivacaine and HOF-TATB@lidocaine HCl significantly suppressed TC severity, with the lidocaine group exhibiting near-complete behavioral rescue over the 6-week observation window (Fig. 5d).

**Fig. 5.**
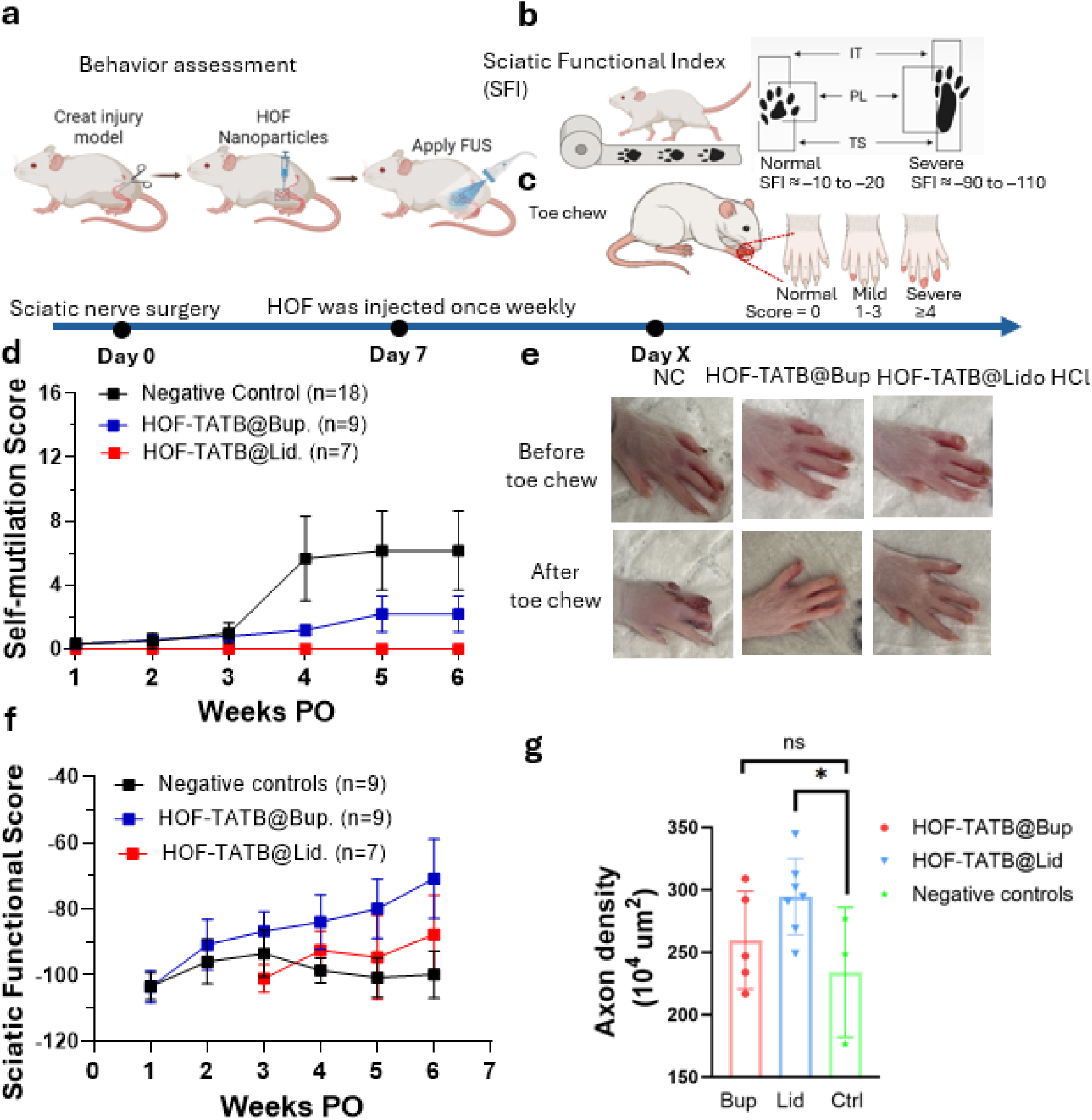
Ultrasound-responsive HOF-TATB nanocarriers mitigate neuropathic pain and enhance motor recovery in a sciatic nerve transection model. (a) Schematic of the rat sciatic nerve transection and repair procedure, nanoparticle treatment, and behavioral assessments. Sciatic nerves were surgically transected and reconnected via microsurgical neurorrhaphy. At post-operative day 7, injured rats received either no treatment (negative control) or local analgesia-loaded HOF-TATB nanoparticles (loaded with lidocaine HCl and bupivacaine) administered to the sciatic nerve region, guided by ultrasound. FUS was then applied to trigger local drug release. From week 2 to week 5, HOF-TATB nanoparticles were injected once weekly and FUS was applied daily. (b,c) Behavioral assessments included the sciatic functional index (SFI) test and evaluation of toe-chewing behavior. (d) Self-mutilation scores in rats receiving HOF-TATB loaded with bupivacaine (blue, n = 9) or lidocaine HCl (red, n = 7), compared to negative Controls (black, n = 18). Both treatment groups showed markedly reduced self-injury behavior over 6 weeks PO, indicating effective attenuation of neuropathic pain by locally released lidocaine and bupivacaine. (e) Representative top-view images of rat paws before and after toe chewing. Negative controls showed severe injury, whereas the HOF-TATB@lidocaine HCl group had minimal or no damage. (f) SFI scores over 6 weeks PO revealed an improved locomotor recovery in rats treated with HOF-TATB@bupivacaine (blue) relative to both NC and the lidocaine-treated group. HOF-TATB@lidocaine HCl (red) showed limited improvement in functional outcomes. (g) Axon density was highest in the HOF-TATB@lidocaine HCl group, significantly exceeding that of negative controls (*p* < 0.05). All data represent mean ± s.d.; statistical analyses were performed using two-way ANOVA with Tukey’s multiple comparisons test unless otherwise noted.

As for motor recovery, as expected, animals receiving neurorrhaphy alone (negative controls) showed minimal improvement in SFI scores over the 6-week period, consistent with previous reports that spontaneous functional recovery is rare in this model. Animals treated with HOF-TATB@bupivacaine demonstrated marked and sustained improvements in SFI scores over time, indicative of functional reinnervation and recovery of locomotor control (Fig. 5f). Moderate gains were also observed in the HOF-TATB@lidocaine HCl treated group, whereas untreated controls showed negligible recovery. The concurrent restoration of both primarily sensory (TC)^15^ and motor primarily (SFI)^51^ functions strongly suggests that ultrasound-programmed anesthetic delivery not only relieves neuropathic pain but also facilitates neurofunctional rehabilitation in the chronic phase of injury. This result might be due to the minimization of aberrant afferent activity and inflammation at the nerve repair site, thereby enabling more favorable conditions for axonal regeneration and functional reintegration. Taken together, these findings position our HOF-based delivery system as a unique neuromodulatory platform—capable of non-invasively reshaping pathological nerve activity, restoring behavioral output, and offering a programmable alternative to current chronic pain and neurorehabilitation therapies. Moreover, HOF-TATB@Lidocaine HCl produced the highest axon density, significantly greater than negative controls (*p* < 0.05). HOF-TATB@Bupivacaine showed a similar upward trend but was not significantly different from controls (Fig. 5e).

This sustained suppression of maladaptive pain TC behavior according to von Frey tests and motor sensory recovery behavior as assessed by the SFI underscores the HOF-TATB platform’s ability to durably modulate aberrant neural activity in a noninvasive, programmable manner. Importantly, these results not only highlight the analgesic efficacy of the HOF drug delivery system under chronic pathological conditions but also reveal its potential as a clinically translatable strategy for long-term neuromodulation following peripheral nerve trauma or repair.

## Discussion

Our *in vivo* behavioral results—derived from machine learning–guided drug selection— provide critical evidence for the translational potential of the HOF-TATB–based ultrasound-responsive drug delivery system. Both lidocaine HCl and bupivacaine were prioritized by our predictive model for their high loading capacity and favorable release profiles, and the *in vivo* testing confirmed their strong analgesic efficacy. Specifically, von Frey testing demonstrated robust, repeatable, and on-demand analgesia, confirming that ultrasound activation can reliably suppress neuropathic pain hypersensitivity in a targeted and temporally precise manner. TC behavior was sustainably suppressed over the multi-week observation period, indicating that the platform effectively mitigates aberrant sensory signaling beyond transient pain relief. Significant improvements in the SFI scores revealed concurrent restoration of voluntary motor function, suggesting that alleviating neuropathic pain not only improves quality of life but also facilitates neurofunctional rehabilitation by reducing maladaptive afferent input, dampening local inflammation, and creating a permissive environment for axonal regeneration. These combined nocioceptive, sensory and motor outcomes—achieved without systemic drug exposure—directly address key clinical requirements for post-operative and chronic pain management. Such outcomes are rarely achieved in current pain control strategies. Importantly, these behavioral results validate the predictive accuracy of our machine learning–guided drug selection, as both model-prioritized candidates demonstrated the high loading, favorable release, and in vivo efficacy predicted by the algorithm. This AI-assisted screening framework could be further extended to accelerate drug–carrier pairing for a wide range of therapeutic agents.

Nevertheless, some limitations remain. Although both lidocaine HCl and bupivacaine showed strong behavioral efficacy, further studies in larger preclinical models are needed to assess long-term stability, biodistribution, and optimal dosing. Future work will explore PEGylation and other rational HOF modifications to enhance drug loading, extend in vivo persistence, and ultimately enabling personalized, programmable neuromodulation for diverse peripheral nerve injuries and chronic pain conditions.

In comparison to the current anesthetic delivery platforms—including liposomes (e.g., EXPAREL), hydrogels, micelles, and polymeric depots—address specific challenges in sustained pain management, but none simultaneously achieve (1) long-term drug retention, (2) high loading efficiency, (3) non-invasive reactivation, and (4) precise spatiotemporal control. Most rely on passive release mechanisms, lack modular programmability, or require surgical implantation, limiting adaptability for chronic or activity-dependent pain. By contrast, our ultrasound-triggered HOF nanoparticle platform integrates all four features into a single injectable system: exceptional drug-loading (∼30.0 wt%) via tailored hydrogen-bonding and π–π stacking, robust *in vitro* retention in serum-rich environments (10% FBS) for over a month, and sustained *in vivo* retention for at least one week, re-triggerable release under clinically safe ultrasound, and modular adaptability for both short- and long-acting anesthetics. This capability enables dynamic responsiveness to change pain states without invasive re-administration and elevates the system from a static drug depot to a programmable neuromodulation interface.

In summary, our programmable system bridges a critical gap in neuromodulation and pain management by enabling non-invasive, externally controlled therapy with minimal systemic exposure. Beyond anesthetic delivery, the modularity of the HOF framework suggests broad applicability for other neurological, inflammatory, and metabolic disorders that require precise spatiotemporal drug control, including epilepsy, post-stroke spasticity, and localized inflammatory pain. Given that the ultrasound parameters used are within clinically approved safety limits and compatible with existing imaging platforms, this approach could be rapidly adapted for translational and clinical deployment. As such, this work lays the foundation for next-generation, ultrasound-guided theranostic platforms in both clinical and translational settings.

## Methods

### HOF-TATB nanoparticles preparation

HOF-TATB nanoparticles were synthesized following a previously reported method with slight modifications. Briefly, 30 mg of H TATB was dissolved in 3 mL of dimethylformamide (DMF) under constant stirring at 1,000 rpm. Subsequently, 12 mL of distilled water was added dropwise, and the mixture was stirred for an additional 10 minutes. The resulting precipitate was collected by centrifugation at 12,000 rpm (13,523 × g) for 5 minutes using an Eppendorf Centrifuge 5420. The collected precipitate was washed three times with acetone and distilled water to remove residual impurities. The final product, with an approximate yield of 30%, was dispersed in water at the desired concentration. The concentration of HOF-TATB in the dispersion was determined using a UV-Vis calibration curve of the H TATB solution.

### HOF-BTB nanoparticles preparation

HOF-BTB nanoparticles were synthesized using a modified procedure. Specifically, 30 mg of H BTB was dissolved in 2 mL of dimethylformamide (DMF) under continuous stirring. Subsequently, 12 mL of distilled water was added dropwise, and the mixture was stirred for an additional 5 minutes. The resulting precipitate was collected by centrifugation at 12,000 rpm (13,523 × g) for 5 minutes using an Eppendorf Centrifuge 5420. The collected nanoparticles were washed three times with methanol and distilled water to remove any residual impurities. The final yield of HOF-BTB nanoparticles was approximately 25%.

### HOF-101 nanoparticles preparation

HOF-101 nanoparticles were synthesized following a modified procedure. Specifically, 30 mg of H TBAPy was dissolved in 3 mL of dimethylformamide (DMF) under constant stirring at 1,000 rpm. Subsequently, 12 mL of distilled water was added dropwise to the solution, and the mixture was stirred for an additional 5 minutes. The resulting precipitate was collected by centrifugation at 12,000 rpm (13,523 × g) for 5 minutes using an Eppendorf Centrifuge 5420. The collected nanocrystals were washed three times with acetone, ethanol, and distilled water to remove any residual impurities. The final product, with a yield of approximately 95%, was resuspended in distilled water at the desired concentration for future use.

### HOF-102 nanoparticles preparation

HOF-102 nanoparticles were synthesized using a modified procedure. Specifically, 10 mg of H PTTNA monomer was dissolved in 2 mL of dimethylformamide (DMF) under constant stirring. Subsequently, 8 mL of methanol was added, and the mixture was stirred for an additional 5 minutes. The resulting precipitate was collected by centrifugation at 12,000 rpm (13,523 × g) for 5 minutes using an Eppendorf Centrifuge 5420. The collected nanocrystals were washed three times with methanol and distilled water to remove any residual impurities. The final product exhibited a yield of approximately 85%.

### Preparation of hydrophilic lidocaine HCl or bupivacaine-HCl-loaded HOF nanoparticles

Lidocaine HCl or bupivacaine HCl-loaded HOF nanoparticles were prepared using a solution immersion method. Briefly, 3 mg of lidocaine HCl or bupivacaine HCl was dissolved in 2 mL of a 5 mg/mL HOF-TATB or other HOF nanocrystal suspension. The mixture was incubated under gentle vibration at 30°C overnight to facilitate drug loading. Following incubation, the nanocrystals were collected by centrifugation at 12,000 rpm (13,523 × g) for 5 minutes using an Eppendorf Centrifuge 5420. The resulting pellets were washed three times with distilled water to remove any unloaded drug molecules. A 1 mL suspension of the drug-loaded nanocrystals was subjected to freeze-drying to obtain a dry powder. Subsequently, 1 mg of the dried powder was dissolved in 0.2 mL of a DMSO and methanol mixture to completely release the loaded drug and TATB ligands. The drug-loading content was determined using high-performance liquid chromatography (HPLC).

### Preparation of lipophilic lidocaine or bupivacaine-loaded HOF nanoparticles

Lipophilic lidocaine or bupivacaine-loaded HOF nanoparticles were prepared using a dual-solvent method. Specifically, 4 mg of lidocaine or bupivacaine was dissolved in 100 µL of dichloromethane (CH Cl ). This solution was then added dropwise to 2 mL of a 10 mg/mL HOF-TATB or other HOF nanoparticle suspension under continuous stirring at 700 rpm for 4 hours to facilitate drug loading. Following the loading process, the nanoparticles were collected by centrifugation at 12,000 rpm (13,523 × g) for 5 minutes using an Eppendorf Centrifuge 5420. The resulting pellets were washed three times with acetone and distilled water to remove any unloaded drug molecules. A 1 mL suspension of the drug-loaded nanoparticles was subsequently subjected to freeze-drying to obtain a dry powder. For drug content determination, 1 mg of the dried powder was dissolved in 0.2 mL of a DMSO and methanol mixture, ensuring complete release of the loaded drug and the amount of the TATB ligands. The drug-loading content was quantified using HPLC.

### Computational Methods

The interaction energy between a drug molecule and a HOF channel was computed the following way with GFN2-xTB^2^. The HOF channel was extracted from the crystal structure and relaxed by restrained minimization. The drug was then placed in the middle of the channel and minimized at the presence of the channel while the channel is completely fixed during the process, which provided the total energy of the drug in the complex. The total energy of the drug minimized without the complex was also obtained. The difference is calculated as the interaction energy. All calculations were done with implicit solvation of water.

### *In vitro* ultrasound-controlled drug uncaging

The *in vitro* release of drugs from HOF nanoparticles under ultrasound stimulation was evaluated using a FUS system. Freshly prepared drug-loaded HOF nanoparticles were suspended in distilled water (pH adjusted to 2) at a concentration of 1 mg/mL and transferred into glass vials. These vials were placed directly on the ultrasound transducer, and the suspension was subjected to FUS irradiation at a frequency of 1.5 MHz under the specified parameters and for predetermined durations. At designated time intervals, 100 µL of the suspension was carefully extracted from the vial and centrifuged at 8,000 rpm (6,010 × g) for 5 minutes using an Eppendorf Centrifuge 5420. The released drug was detected in the supernatant, and its concentration was determined using HPLC. The percentage of drug release was calculated based on the initial drug loading content.

### Long-term drug stability evaluation in HOF nanoparticles

The long-term stability of drug-loaded HOF nanoparticles was evaluated under physiological conditions. Drug-loaded HOF nanoparticles were prepared at a concentration of 3 mg/mL in fetal bovine serum (FBS) solutions of either 10% or 50% and stored in glass vials at room temperature. At predetermined time intervals, 100 µL of the nanoparticle suspension was withdrawn and centrifuged at 8,000 rpm (6,010 × g) for 5 minutes using an Eppendorf Centrifuge 5420. The collected nanoparticles were washed three times with acetone and distilled water (DIW) to remove any loosely bound drug. A 1 mL suspension of the washed nanoparticles was then subjected to freeze-drying, resulting in a dry powder. For drug content determination, 1 mg of the dried powder was dissolved in 0.2 mL of a dimethyl sulfoxide (DMSO) and methanol mixture, ensuring complete release of the loaded drug and dissociation of the TATB ligands. The drug-loading content was quantified using HPLC.

### *In vitro* calcium imaging

Calcium imaging was performed using primary cortical neurons transduced with pAAV(adeno-associated virus)-hSyn-GCaMP6s-WPRE-SV40. Following a 6-day transduction period, the neuron cultures were placed on the stage of a Leica DMi8 fluorescence microscope equipped with a 20× air objective. The water balloon membrane of the FUS transducer was brought into direct contact with the neuron culture medium atop the plate. Fresh TATB@Lido nanoparticles were added to the medium at a final concentration of 5 µg/mL Lido. FUS stimulation (1.08 MPa, 1.40 MHz, 10 s duration) was applied to trigger the release of Lido, leading to neuronal inhibition. Fluorescence imaging videos were captured using the Leica DMi8 microscope under the green fluorescence channel (Ex: 450–490 nm) with a 5 ms exposure time. For data analysis, the transient increase in green fluorescence (ΔF/F) was determined by manually segmenting 50 neurons and extracting the fluorescence time-series data. The imaging videos were converted to grayscale using ImageJ software, and the fluorescence data were processed using a custom MATLAB algorithm. This algorithm performed detrending and normalization of the fluorescence time-series data through second-order polynomial curve fitting and baseline maximum fluorescence value extraction, effectively compensating for photobleaching effects.

### Animal experiments

All animal studies were conducted in accordance with the National Institutes of Health Guide for the Care and Use of Laboratory Animals (8th ed., National Research Council, 2011) and approved by the Institutional Animal Care and Use Committee (IACUC) at the University of Texas at Austin (AUP-2022-00278, approved 02/07/2023). Male outbred Sprague-Dawley (SD) rats aged 3–6 months were housed 2–3/cage under a 12-hour light/dark cycle. Food and water were provided ad libitum. Surgical and behavioral procedures were conducted during the day. The rats were randomly assigned to each group. Under brief isoflurane–oxygen anesthesia, operated (see below in Surgical Procedures) or unoperated (for in vivo nanoparticle evaluation) rats were injected with 500μL of HOF-TATB with or without lidocaine HCL or bupivacaine using a 23G needle. The injection was performed next to the sciatic nerve with the needle introduced posteromedial to the greater trochanter. Ten minutes after the injection, focused ultrasound activation was performed at the injection site.

### Ultrasound-triggered sciatic nerve blockade in unoperated Sprague Dawley rats

Following local injection of nanoparticles (either unloaded, HOF-TATB@Lidocaine·HCl, or HOF-TATB@Bupivacaine) at the sciatic nerve, focused ultrasound (FUS; 1.5 MHz, 1.40 MPa) was applied with a pulsed sequence of 0.2 s on / 0.8 s off for the designated activation period. 30 minutes after ultrasound stimulation, von Frey (VF) testing was performed under anesthesia to assess nerve blockade. Mechanical nociceptive thresholds were determined by measuring paw withdrawal responses to VF filaments applied to the dorsal region of the hindpaw between digits 4 and 5.

### Sensory and motor behavioral testing

Dorsal sensory von Frey (VF) testing was performed as previously described^52,53^. Briefly, prior to the administration of drugs (unoperated group) or surgery (peripheral nerve injury group), rats were handled and trained daily for at least 4 sessions. VF testing was performed at predetermined time points for unoperated rats or weekly time points for operated rats. The second tester applied the VF filaments when the rat was calm. The first tester gently wrapped and restrained rats in a towel and elevated them to an upright position to expose the dorsal surface of their hindpaws. The second tester applied VF filaments (Stoelting Co., Wood Dale, IL, USA) between digits 4 and 5 when rats were calm. The beginning filament was presented verticallyperpendicular to the targeted dorsal region until bent for two1-2 seconds and was repeated on the alternate paw. A positive or negative withdrawal response was recorded for each trial on each side hindpaws. A positive and negative withdrawal responses were respectively followed by a lower and higher force filament on the subsequent trial. Five trials were tested on each side hind paw. Rats were allowed to rest at least 30 seconds between trials of both hindpaws. The final VF thresholds depended on the last response obtained and were converted to gram force for analysis. Any response was voided and the trial was repeated if the filament slipped off of the dorsal surface, the rat moved or picked up its hind paw, or if the rat struggled in the towel during application of the filament. VF filaments were routinely tested on a scale measuring grams and replaced to accurately represent intended gram forces.

Sciatic function index (SFI) testing was performed as previously described. All rats were trained daily for three days prior to surgery. SFI testing was conducted weekly in all rats until their experimental PO endpoints. SFI testing and grading was completed with testers blinded to the experimental groups. Rat hindpaws were inked blue on the operated side and red on the unoperated side. Rats were placed on a slightly inclined 100 mm-wide wooden board lined with white paper and allowed to return to their home cage. Two trial runs without stopping or hesitation were gathered for each testing day. A successful SFI trial included three consecutive steps on each hind paw without stopping or hesitating. Each rat completed two successful trial runs were gathered for each on their testing day. An average score was calculated for that day using the following formula: 73*((NPL-EPL)/EPL+(ETS-NTS)/NTS+(EIT-NIT)/NIT, where NPL stands for normal footprint length, EPL for experimental footprint length, NTS for normal toe spread, ETS for experimental toe spread, NIT for normal intermediary toe spread, and EIT for experimental intermediary toe spread.

### Surgical procedures

Surgical procedures were performed as previously described^52,53^. Rats were anesthetized with isoflurane (4% induction, 2% maintenance (RXISO-250; Animal Health International, Roanoke, TX, USA))/oxygen mixture at 1.5 L/min (Handlebar Anesthesia, Pflugerville, TX, USA). The lateral aspect of the left hindlimb was shaved and sterilized, and a 1.5-2 cm incision was made through the skin and the biceps femoris to expose the sciatic nerve. The sciatic nerve was sharply transected at the mid-thigh level with microscissors. Sciatic nerve stumps were irrigated with 0.5% methylene blue followed by a hypotonic (250 mOsm), calcium-free diluted Normosol-R (ICU Medical, San Clemente, CA, USA) saline solution. Sciatic nerve stumps were trimmed, closely apposed, and secured with at least four 10-0 microsutures (neurorrhaphy) through the epineurium sheaths.

Following neurorrhaphy, lesion sites were flushed several times with Ca^2+^-containing normal saline. The surgical site was closed with 5-0 sutures through the muscle, and the skin was closed with wound clips. Rats were allowed to recover on heat pads before being returned to standard housing. Rats received 5 mg/kg subcutaneous injections of carprofen during surgery and for three days PO.

### Self-mutilation scoring

Self-mutilation often first presented as nibbling of the most distal parts of the toe nails with continued chewing in the proximal direction of the hind paw digits. The severity and the time of onset of self-mutilation to the operated hind limb were monitored and recorded daily until the PO endpoint was reached by either euthanasia criteria set by the IACUC or by reaching an experimental endpoint. Self-mutilation severity ranged from nail bed bleeding to bone exposure and received different scores. Animals received a score of 1 for each digit with significant tissue damage that required additional PO anti-inflammatory carprofen administration, as determined by veterinarian staff. An additional score of 5 was given when any digit had bone exposure. For example, when 3 digits had tissue damage and one digit had bone exposure, the rat received a score of 8; two digits had tissue damage and 2 digits had bone exposure, the rat received a score of 12. Self-mutilation rates are presented as percentages and scores are presented as mean ± SEM.

### Morphological analyses for the sciatic nerve and the soleus muscle

After rats were deeply induced with 4% isoflurane/oxygen euthanized by an intracardiac KCl injection, sciatic nerves and soleus muscles were harvested for following analyses.

Nerve morphometric analysis: sciatic nerves harvested from operated limbs were placed in 0.1 M sodium cacodylate buffer and then fixed by 2% paraformaldehyde/3% glutaraldehyde fixative overnight. The next day, tissues were washed with buffer, trimmed and postfixed in 1% osmium tetroxide/1% potassium ferrocyanide in 0.1 M sodium cacodylate buffer for 3 to 5 hours, washed in water, stained in 1% aqueous uranyl acetate for 1 to 2 hours, and then washed and held in water^54^. Nerves were dehydrated using graded alcohols, exchanged to absolute acetone, placed in increasing concentrations of Hard Plus Resin 812 (Electron Microscopy Sciences, Hatfield, PA), and then embedded in fresh resin and polymerized at 60°C. Glass knife thick sections (0.5 mm) were stained in toluidine blue and imaged using confocal microscopy. Axon numbers per square micrometer (mm^2^) were counted and averaged from at least 3 different regions of interest containing at least 150 axons per sample.

Soleus muscle isolation: Soleus muscles from both unoperated and operated limbs were weighed immediately after isolation. The muscle weight ratio was calculated by dividing the weight of the operated side by that of the unoperated side.

### Ultrasound-triggered pain management and motor functional recovery after sciatic nerve injury (SNI)

Self-mutilation (autotomy) of the hind limbs in SD rats following SNI is a well-established model for studying neuropathic pain^12,14,16^. In this model, rats underwent unilateral sciatic nerve transection followed by end-to-end microsuture repair under an operating microscope. Post-surgery, animals were monitored daily for signs of distress and self-mutilation scores were recorded until 6 weeks post-operation (PO). To evaluate the therapeutic effects of HOF-TATB@lidocaine and HOF-TATB@bupivacaine, nanoparticles were perineurally injected at the repair site, and focused ultrasound stimulation (1.5 MHz, 1.40 MPa, 8 min) was applied daily to trigger on-demand anesthetic release. Motor functional recovery was assessed weekly for 6 weeks PO using the SFI.

### Mass Spectrometry Imaging

Frozen nerve specimens were longitudinally sectioned at 12 µm thickness using a Thermo NX50 CryoStar cryostat and collected onto ColorFrost Plus™ slides. Serial sections were collected for H&E staining. Fiducial points were placed at the corners of the slides for MSI prior to collection of an optical image using an Epson Perfection V600 flatbed document scanner at 4800 dpi. The sections were coated with 40 mg/mL 2,5-dihydroxybenzoic acid in 50% acetonitrile, 0.1% trifluoracetic acid using an HTX M5 Robotic Reagent Sprayer over 10 passes with the following parameters: a flow rate of 100 µL/min, a track speed of 1200 mm/min, a track spacing of 2 mm, a nozzle temperature of 75°C, a nozzle height of 40 mm, a heated tray temperature of 50°C, a nitrogen pressure of 10 psi, and a CC track pattern. Mass spectrometry image data were acquired at 30 µm resolution using FlexImaging 7.0 on a Bruker timsTOF fleX QTOF mass spectrometer in positive ion mode with the following parameters: 300 shots per pixel, an *m/z* range of 50-1000, a Funnel 1 RF of 150.0 Vpp, a Funnel 2 RF of 200.0 Vpp, a Multipole RF of 200.0 Vpp, a Collision Energy of 5.0 eV, a Collision RF of 700.0 Vpp, a Transfer Time of 80.0 µs, and a Pre Pulse Storage of 8.0 µs. MSI data was loaded into SciLS Lab 2025b and normalized to Total Ion Current for visualization of the lidocaine distribution in the nerve sections.

### Statistical analyses

Excel was used to calculate means and standard deviations. Graphpad Prism (version 9, GraphPad Software, Boston, MA, USA, www.graphpad.com) was used to perform linear regressions and t-test comparisons.

## Acknowledgment

TEM image acquisition was performed with the help of Michelle Mikesh at the Center for Biomedical Research Support Microscopy and Imaging Facility at UT Austin (RRID# SCR_021756). Prof. Huiliang Wang acknowledges funding support from the National Science Foundation (NSF) CAREER award (2340964), NIH Maximizing Investigators’ Research Award (National Institute of General Medical Sciences 1R35GM147408), UT Austin Portugal Exploratory Research Projects Grant, Robert A. Welch Foundation Grant (No. F-2084-20240404) and Craig H. Neilsen Foundation Pilot Research Grant. We acknowledge BioRender.com for the figures drawing.

## Author Contributions

W.H., C.Y., X. S., Y.W. P.R., G.B. and H.W. designed the project. W.H. led the materials design and characterization. W.H., C.Y., X. S., B. A., L. Z., H.G., A.S. and A.O. conducted the animal tests. W.H., W.W., X.L., K.W.K.T., J.J., S.Y., A.R.L. conducted in cell culture test and analysis. Y.W., Z.H. developed the machine learning model for drug screening. E.H.S. conducted MSI analysis. All the authors contributed to the writing of the manuscript.

## Competing Interests Statement

The authors H.W., W.H., C.Y., X. S., Y.W., W.W., P.R., and G. B. declare that a patent application (UTSB.P1394US.P1) relating to this work has been filed. The other authors declare no competing interests.

## Additional Information

Supplementary Information is available for this paper.

## Notes

### Competing Interest Statement

The authors declare that a provisional patent application (UTSB.P1394US.P1) relating to this work has been filed. The other authors declare no competing interests.

