## Supplementary Information for "A machine-learning-guided hydrogen-bonded organic framework for long-term, ultrasound-triggered pain therapy"

**Supplementary Information Guide:**

**Supplementary Methods** (pages 3-7)

1. Materials and Instrumentation. (page 3)
2. Preparation of different drug-loaded HOFs and evaluation of ultrasound-triggered drug release through high-performance liquid chromatography (HPLC). (page 4-5)
3. Primary neuron culture. (page 5)
4. Ultrasound-triggered sciatic nerve blockade in unoperated Sprague Dawley rats. (page 5-6)
5. Iba and Caspase-3 Staining. (page 6)
6. H&E staining. (page 6)
7. Computational Methods. (page 6-7)

**Supplementary Figs 1-10.** (page 8-17)

**Supplementary Tables 1-2.** (page 18-25)

**Supplementary References.** (page 25)

***Materials and Instrumentation***

***Chemicals.*** Unless otherwise specified, all reagents and solvents were obtained from commercial suppliers and used without further purification. The following chemicals were used in this work: concentrated hydrochloric acid (HCl), acetone, dimethylformamide (DMF), dichloromethane (DCM), methanol, ethanol, and dimethyl sulfoxide (DMSO). 1,3,5-Tris(4-carboxyphenyl)benzene ($H_3BTB$), 4,4′,4″-(1,3,5-triazine-2,4,6-triyl)tribenzoic acid ($H_3TATB$), 1,3,6,8-tetrakis(benzoic acid)pyrene ($H_4TBAPy$), and 1,3,6,8-tetra(6-carboxynaphthalen-2-yl)pyrene ($H_4PTTNA$) were purchased from Chemscene.

**Instruments**.

Transmission electron microscopy (TEM) images of the HOFs were acquired using a JEOL NEOARM low-kV, aberration-corrected TEM/STEM (30–200 kV) at the Texas Materials Institute, The University of Texas at Austin, in TEM mode at an accelerating voltage of 200 kV. The powder X-ray diffraction (PXRD) patterns were collected using a Rigaku Miniflex 600 diffractometer equipped with a Cu Kα radiation source (λ = 1.54184 Å) operating at 40 kV and 15 mA. Data were recorded over a 2θ range of 2.5–30°. An Agilent MR400 spectrometer ($^1$H, 400 MHz) was used for the collection of $^1$H NMR spectra. UV–Vis spectra were recorded on an Eppendorf BioSpectrometer® Basic. The drug release percentage from HOFs was measured using HPLC (Agilent 6120 Single Quadrupole LC/MS).

**Bioreagents**. AAV-hSyn-GCaMP6s-WPRE-SV40 (Addgene viral prep #100843-AAV9; RRID:Addgene_100843) was obtained from the Douglas Kim & GENIE Project (http://n2t.net/addgene:100843)[1]. Lidocaine, lidocaine hydrochloride, bupivacaine, bupivacaine hydrochloride, aspirin, benzocaine, carbamazepine, ilomastat (GM6001), ibuprofen, isradipine, tetracaine, deschloroclozapine (DCZ), (S)-3,5-dihydroxyphenylglycine (DHPG), dopamine, L-dopa, methylene blue, methylprednisolone, rhodamine B, and scopolamine were purchased from Sigma-Aldrich and used without further purification.

**Determination of Drug Loading Content**. The drug loading content was determined either by HPLC or by $^1$H NMR, depending on the analytical suitability of each drug. For HPLC analysis, DMSO was used to dissolve the samples prior to injection, unless otherwise specified. For $^1$H NMR analysis, samples were dissolved in DMSO-$d_6$, and the drug content was calculated based on the integration of characteristic peaks.

**Preparation of different (lipophilic drug) drug-loaded HOF-TATB and evaluation of ultrasound-triggered drug release through high-performance liquid chromatography (HPLC).** To prepare drug-loaded TATB-HOF, 2 mg of drug was dissolved in 2 mL of HOF-TATB suspension (5 mg/mL). The mixtures were incubated at 37 °C for 10 h and centrifuged at 12,000 rpm (13,523 × g) for 5 min. The resulting pellets were washed three times with distilled water to remove unloaded cargoes and then re-suspended. To evaluate the drug-loading content, 1 mL of the suspension was collected and freeze-dried. The resulting powder (~1 mg) was dissolved in 0.2 mL of DMSO to completely release the cargo, followed by dilution with 0.4 mL of methanol. The drug-loading content was measured using HPLC (Agilent 6120 Single Quadrupole LC/MS).

For evaluating the ultrasound-triggered drug release, freshly prepared drug-loaded HOF nanocrystal suspensions (10 mg/mL) were transferred into glass vials and placed on an ultrasound transducer. The samples were exposed to focused ultrasound (FUS) at 0.79 MPa, 1.40 MPa, 2.45 MPa and 1.5 MHz for the designated duration under the specified parameters. At predetermined time points, 100 μL aliquots were withdrawn and centrifuged at 8,000 rpm (6,010 × g) for 5 min. The supernatants containing the released drug were collected, and the release percentage was determined based on HPLC calibration curves.

**Preparation of HOF-BTB loaded with various drugs (lipophilic drug) and analysis of ultrasound-responsive release via HPLC.** HOF-BTB suspensions (5 mg/mL, 2 mL) were combined with 2 mg of the desired drug and gently agitated to ensure uniform mixing. The mixtures were maintained at 37 °C for 10 h, after which they were centrifuged at 12,000 rpm (13,523 × g) for 5 min. The resulting pellets were washed three consecutive times with distilled water to remove excess, non-incorporated drug and then dispersed again in water. For drug-loading measurements, a 1 mL portion of the suspension was collected, freeze-dried, and the obtained powder (~1 mg) was first dissolved in 0.1 mL DMSO to fully release the encapsulated molecules, followed by dilution with 0.5 mL methanol. The final solutions were analyzed using HPLC (Agilent 6120 Single Quadrupole LC/MS) to determine the drug content.

**Preparation of HOF-101 loaded with various drugs (lipophilic drug) and analysis of ultrasound-responsive release via HPLC.** HOF-101 suspensions (5 mg/mL, 2 mL) were mixed with 2 mg of the selected drug under gentle agitation to achieve homogeneous dispersion. The mixtures were incubated at 37 °C for 10 h and then centrifuged at 12,000 rpm (13,523 × g) for 5 min. The pellets obtained were washed three times with distilled water to remove unbound drug and subsequently re-suspended. For drug-loading quantification, 1 mL of the suspension was freeze-dried, and the resulting powder (~1 mg) was dissolved directly in 0.5 mL DMSO to completely release the encapsulated drug. The drug content was determined using HPLC (Agilent 6120 Single Quadrupole LC/MS).

**Preparation of HOF-102 loaded with various drugs (lipophilic drug) and analysis of ultrasound-responsive release via HPLC.** A suspension of HOF-102 (5 mg/mL, 2 mL) was combined with 2 mg of the target drug and stirred gently to ensure even dispersion. The mixture was kept at 37 °C for 10 h, followed by centrifugation at 12,000 rpm (13,523 × g) for 5 min. The precipitates were rinsed three times with distilled water to eliminate residual, unbound drug and then re-dispersed in water. For determining the drug loading, 1 mL of the suspension was taken, freeze-dried, and the obtained solid (~1 mg) was dissolved in 0.5 mL DMSO to fully release the encapsulated molecules. The drug concentration was quantified by HPLC (Agilent 6120 Single Quadrupole LC/MS).

**Primary neuron culture.** C57BL/6 mice (8 weeks old, 20–26 g; Jackson Laboratory) were used in this study. Primary cortical neurons were isolated from embryos at embryonic day 15.5. Briefly, 24-well cell culture plates were coated with poly-L-ornithine (0.2 mg/mL) and incubated at 37 °C for 2 h. The plates were then rinsed three times with PBS and pre-warmed in a cell incubator for 15 min before use. Dissociated neuronal cells were seeded onto the coated plates and maintained in Neurobasal medium supplemented with B27, glutamine, penicillin, and streptomycin. Cultures were incubated at 37 °C in a humidified atmosphere containing 7% $CO_2$. After 2 days in vitro, a glial inhibitor, 5-fluoro-2′-deoxyuridine (0.1 mM), was added to the culture medium. On day 4, 1 μL of pAAV-hSyn-GCaMP6s-WPRE-SV40 was transfected into the neurons. Calcium imaging experiments were performed after an additional 6 days of incubation.

**Ultrasound-triggered sciatic nerve blockade in unoperated Sprague Dawley rats.** After confirming the spatial localization and retention of anesthetic agents in the sciatic nerve of Sprague Dawley (SD) rats as described above, sciatic nerve blockade was evaluated via local administration of anesthetic formulations. Nanoparticles without drug (negative control) or loaded with either HOF-TATB@Lidocaine HCl or HOF-TATB@Bupivacaine were injected locally at the sciatic nerve. This was followed by focused ultrasound (FUS; 1.5 MHz, 1.40 MPa) treatment and von Frey (VF) testing. Blockade of mechanical nociceptive responses was assessed by measuring paw withdrawal thresholds in response to VF filament stimulation applied to the dorsal surface of the hindpaw between digits 4 and 5.

**Iba1 and Caspase-3 Staining.** Following the *in vivo* ultrasound activation procedures, rats were deeply anesthetized with 4% isoflurane in oxygen and euthanized via intracardiac KCl injection. Sciatic nerves were harvested, fixed in 4% paraformaldehyde overnight at 4 °C, and sectioned using a cryostat. Longitudinal sciatic nerve sections (10 μm thickness) were washed with 0.3% TBS and blocked with 5% bovine serum albumin in TBS for 30 min at room temperature. The blocking buffer was then replaced with one of the following primary antibody solutions (prepared in TBS): rabbit anti-Iba1 (013-27691, Wako Chemicals, 1:500), rabbit anti-cleaved Caspase-3 (9661, Cell Signaling Technology, 1:500). After overnight incubation at 4 °C, sections were washed three times with TBS and incubated for 2 h at room temperature in the dark with a secondary antibody solution containing donkey anti-rabbit Alexa Fluor 594 (A32754, Invitrogen, 1:500) and Hoechst 33342 (17535, ATT Bioquest, 1:5000). Finally, sections were washed three times with TBS, mounted on glass slides with mounting medium, and coverslipped. Fluorescence images were acquired using a Nikon AXR-NSPARC confocal fluorescence microscope.

**H&E staining.** Following the *in vivo* ultrasound activation procedures, rats were deeply anesthetized with 4% isoflurane in oxygen and euthanized via intracardiac KCl injection at 14 days post-treatment. Sciatic nerves were harvested, fixed in 4% paraformaldehyde overnight at 4 °C, and sectioned into 10 μm slices using a cryostat. Hematoxylin and eosin (H&E) staining was performed on the sciatic nerve sections according to previously described protocols. The stained sections were mounted on glass slides with mounting medium and coverslipped. Images were acquired using a Nikon Upr-FLM compound light microscope (Nikon Instruments).

**Computational Methods.** Structural descriptors for HOFs and drugs were selected based on the preliminary understanding that non-specific interactions, such as hydrophilicity, play a dominant role in drug loading into HOF crystals. For drugs, the descriptors included molecular volume, radius of gyration, molecular weight, number of rotatable bonds, ratio of $sp^3$-hybridized carbons, total polar surface area, logP, and net charge at pH 7. For HOFs, the descriptors included the number of hydrogen-bonding sites, crystal pore volume, crystal pore size, BET surface area, and logP. All non-crystal features were calculated using RDKit[2].

The interaction energy between a drug molecule and a HOF channel was calculated using the GFN2-xTB method[3]. A HOF channel was extracted from the crystal structure and relaxed via restrained minimization. The drug molecule was positioned at the center of the channel and minimized in the presence of the fixed channel to obtain the total energy of the complex. Separately, the drug molecule was minimized without the HOF channel to obtain its total energy in isolation. The interaction energy was determined as the difference between these two values. All calculations were performed with implicit solvation in water.

### Supplementary Figures

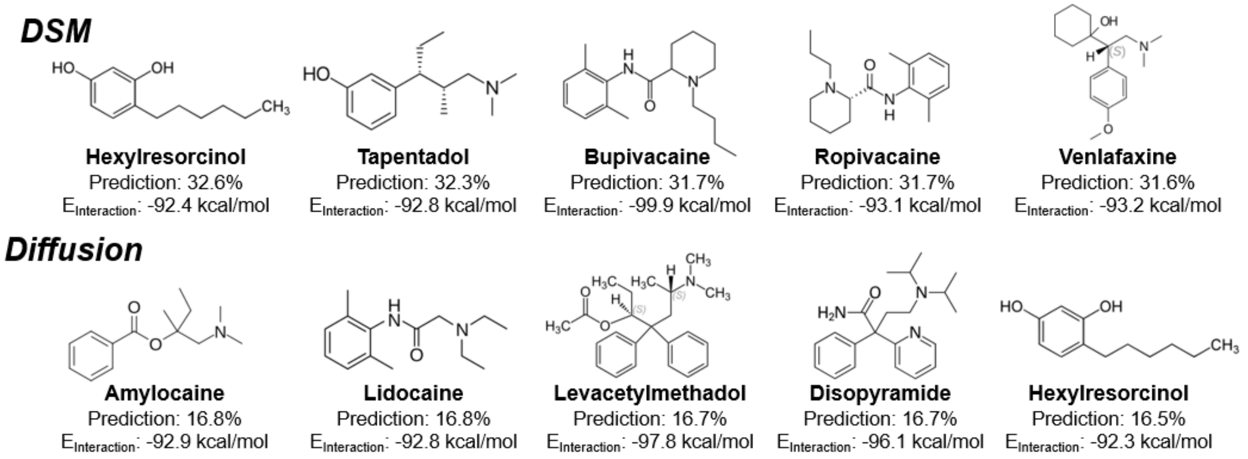


**Supplementary Fig. 1.** Following the machine learning–based screening, the top 5 drug candidates for HOF-TATB were selected for further evaluation using two different encapsulation methods: DSM and diffusion. For each method, the ranking was based on the predicted loading efficiency, and the interaction energies ($E_{\text{interaction}}$) between the drug molecules and HOF-TATB channels were calculated using GFN2-xTB with implicit water solvation. The top 5 drugs identified via DSM included hexylresorcinol, tapentadol, bupivacaine, ropivacaine, venlafaxine, whereas the diffusion method yielded amylocaine, lidocaine, levacetylmethadol, disopyramide, hexylresorcinol.

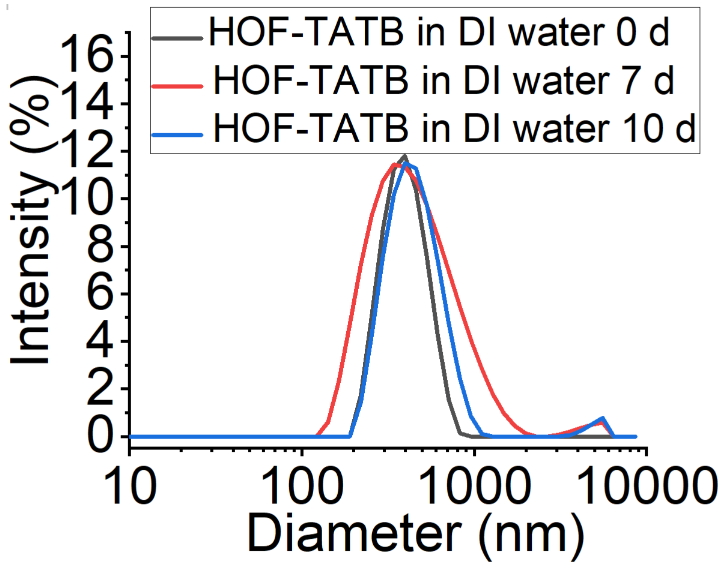


**Supplementary Fig. 2.** Dynamic light scattering (DLS) measurements showed that HOF-TATB nanoparticles maintained a relatively consistent size distribution when dispersed in DI water over a period of 10 days. The initial average hydrodynamic diameter (day 0) exhibited only a slight shift after 7 and 10 days of storage, with no significant aggregation or formation of large particles observed. These results indicate that HOF-TATB possesses good colloidal stability in aqueous environments over extended periods.

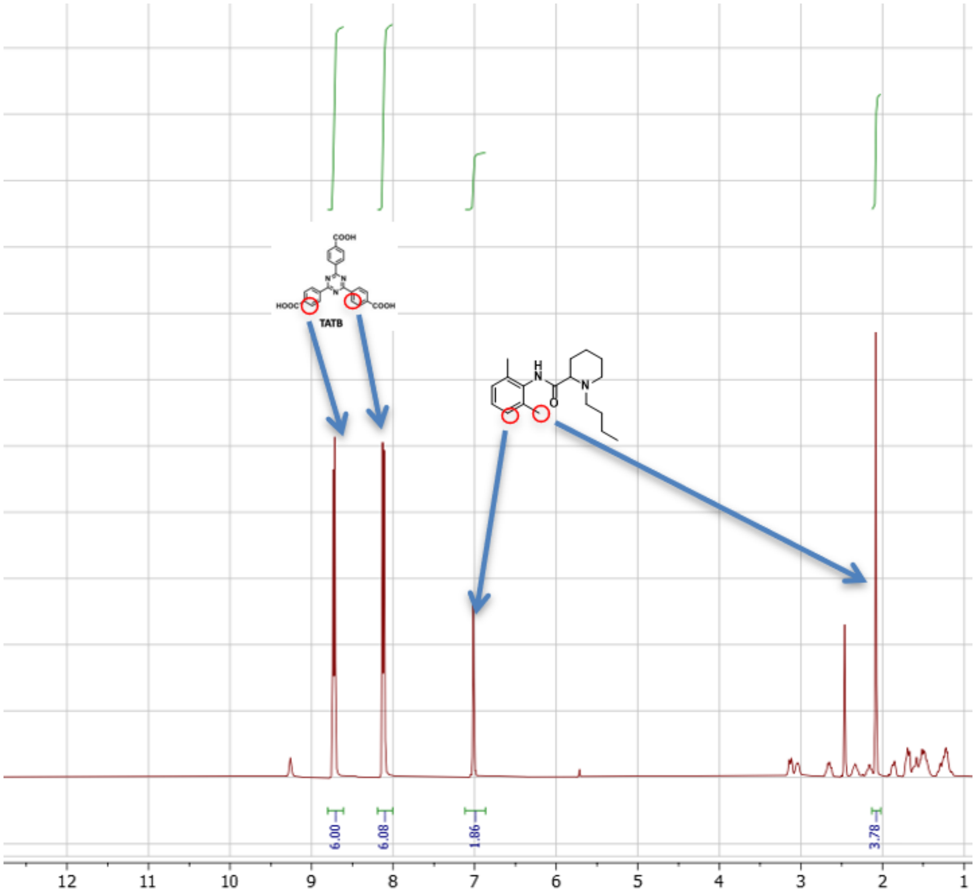


**Supplementary Fig. 3.** $^{1}$H NMR spectrum of HOF–TATB loaded with bupivacaine in DMSO-$d_6$ (400 MHz), showing characteristic peaks of both the HOF–TATB framework and bupivacaine (δ = 7.05 and 2.13 ppm), confirming successful drug loading.

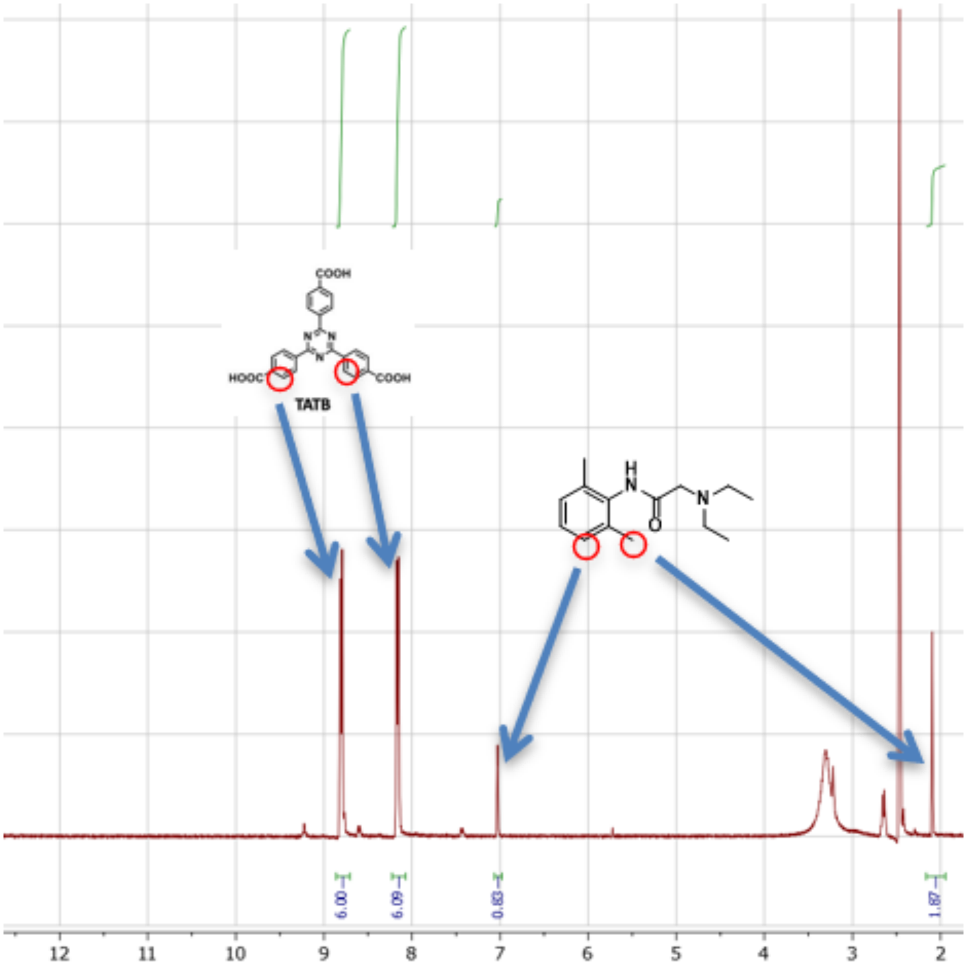


**Supplementary Fig. 4.** $^{1}$H NMR spectrum of HOF–TATB loaded with lidocaine·HCl in DMSO-$d_6$ (400 MHz). Characteristic proton signals from lidocaine·HCl are observed at δ = 7.05 and 2.13 ppm, together with peaks from the HOF–TATB framework, confirming successful drug encapsulation.

.

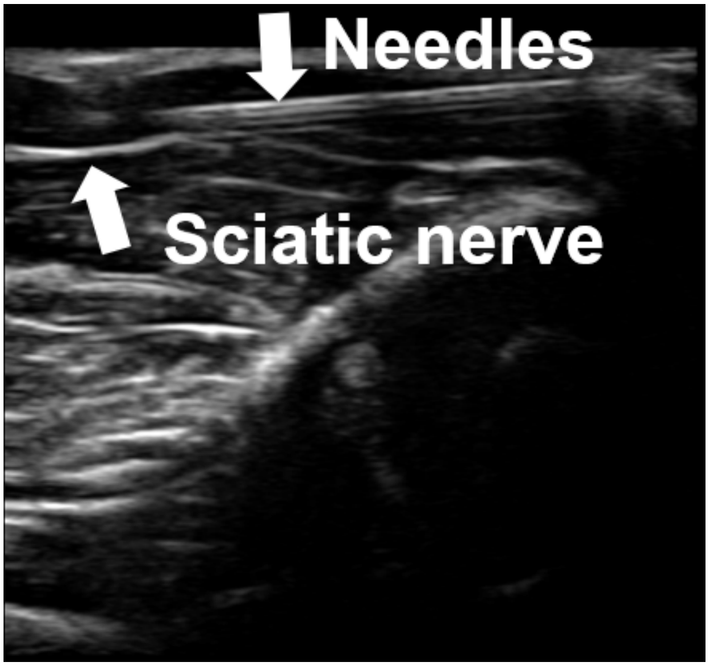


**Supplementary Fig. 5. Ultrasound-guided injection of HOF-TATB nanoparticles.** Representative ultrasound image showing the position of the needle (top arrow) and the sciatic nerve (side arrow) during perineural injection of HOF-TATB nanoparticles. Real-time ultrasound guidance was used to ensure accurate needle placement adjacent to the sciatic nerve for localized delivery of the nanoparticle formulation.

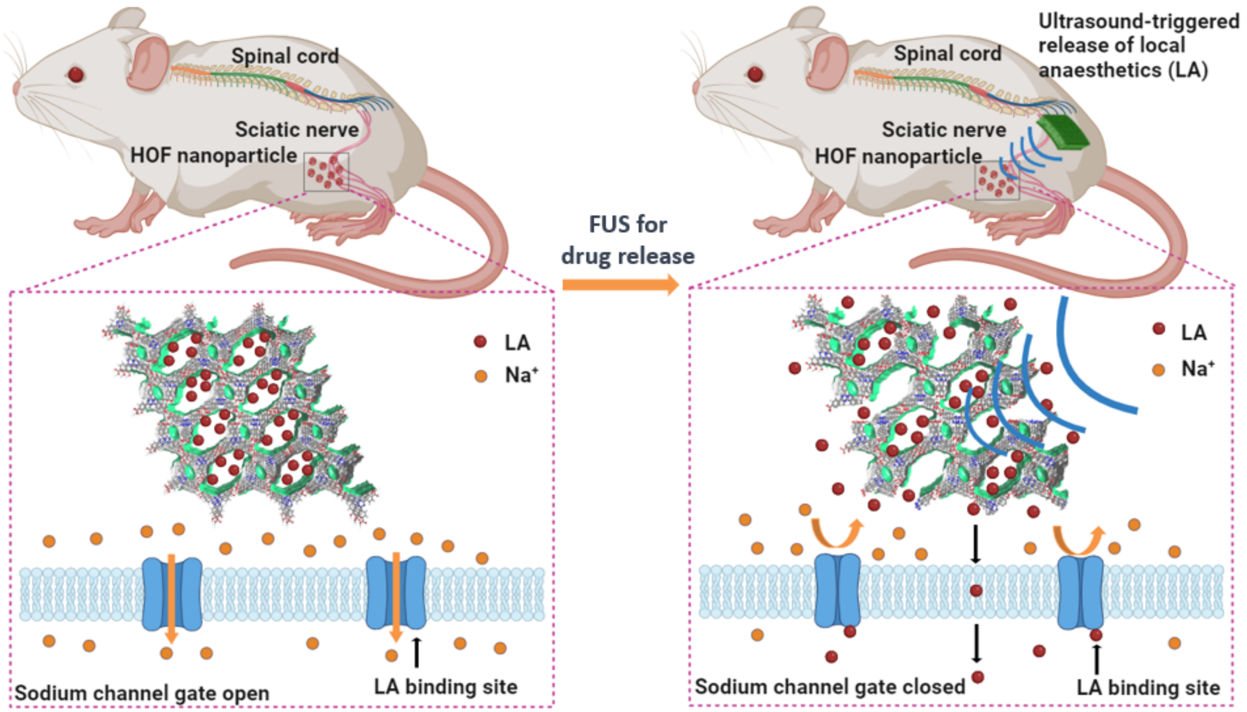


**Supplementary Fig. 6. Schematic illustration of ultrasound-triggered local anaesthetic (LA) release from HOF nanoparticles for sciatic nerve blockade.** HOF nanoparticles loaded with local anaesthetics (LA) were locally administered adjacent to the sciatic nerve. In the absence of focused ultrasound (FUS), LA molecules remain encapsulated within the HOF framework, and sodium channels in the neuronal membrane remain open, allowing $Na^+$ influx and normal nerve conduction. Upon FUS stimulation, the HOF structure releases the encapsulated LA, which binds to sodium channel sites, blocking $Na^+$ influx and thereby inhibiting action potential propagation, resulting in reversible sciatic nerve blockade.

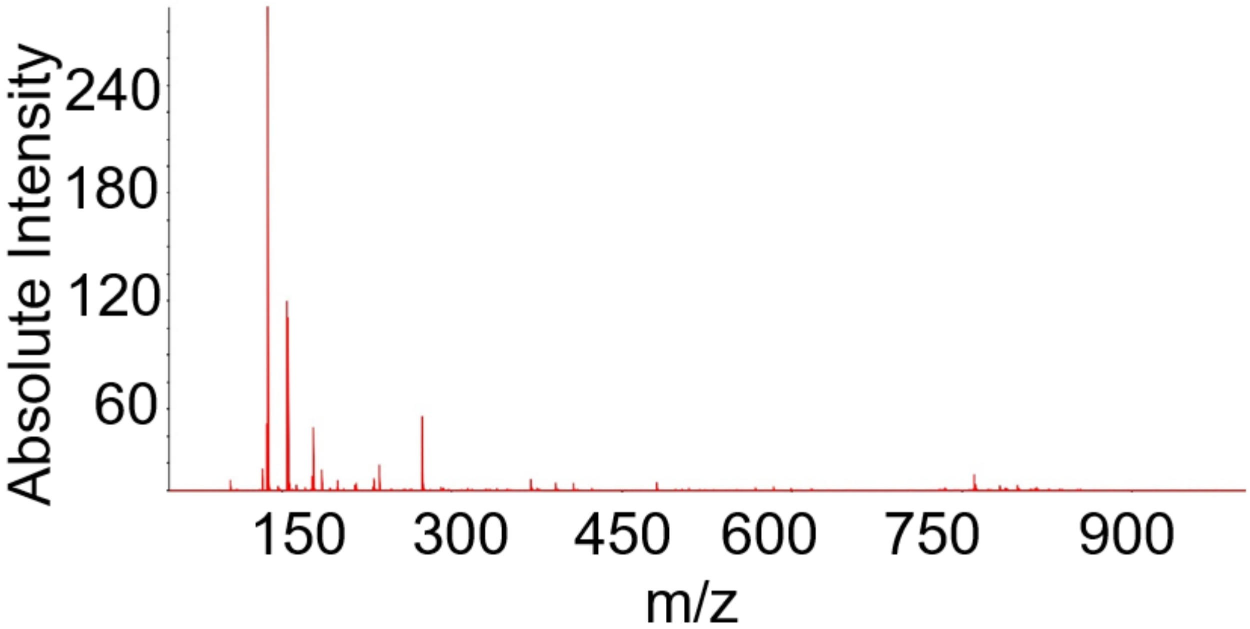


**Supplementary Fig. 7.** Mass spectrometry identification of lidocaine
Mass spectrometry imaging (MSI) analysis revealed a strong signal at *m/z* = 235.1801, which corresponds to the $[M+H]^+$ of lidocaine. This peak, together with its characteristic fragment ions, is consistent with the reference mass spectrum, confirming the presence of lidocaine in the sample.

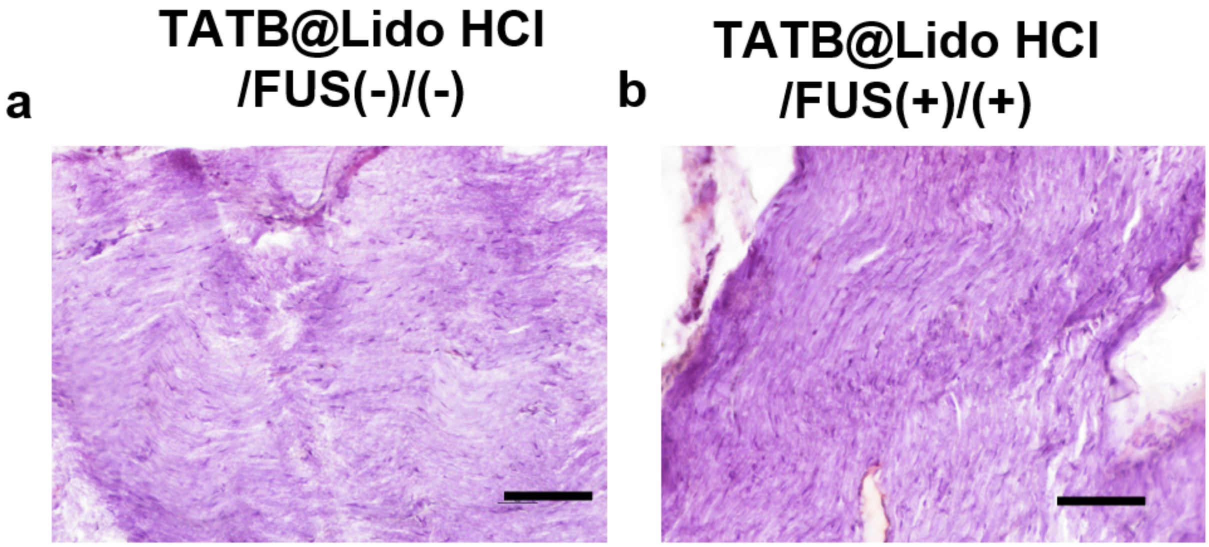


**Supplementary Fig. 8.** Representative hematoxylin and eosin (H&E) staining images of sciatic nerve sections from rats treated with TATB@Lidocaine HCl without focused ultrasound (FUS) activation (a) and with FUS activation (b). For the FUS group, treatment was applied at 1.5 MHz and 1.40 MPa for 8 min. Sciatic nerves were harvested 7 days after treatment. In both conditions, the nerve fibers exhibited normal morphology with intact myelin structure and no signs of degeneration, inflammatory infiltration, or tissue damage, indicating that the application of TATB@Lidocaine HCl, with or without FUS, did not induce noticeable histopathological changes in sciatic nerve tissue (scale bar: 100 μm).

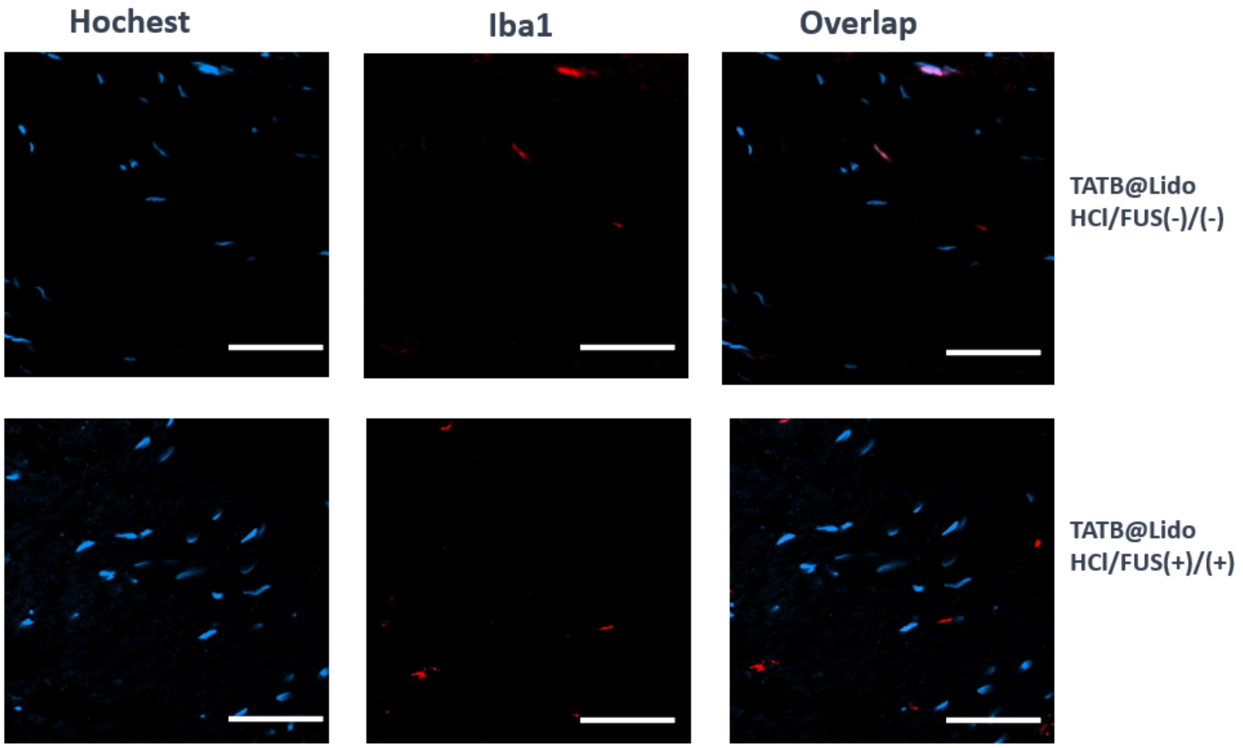


**Supplementary Fig. 9.** Representative immunofluorescence images of rat sciatic nerve sections 7 days after treatment, stained with Hoechst (blue, nuclei) and Iba1 (red, macrophages/microglia). Top: TATB@Lidocaine HCl without FUS activation; Bottom: with FUS activation (1.5 MHz, 1.40 MPa, 8 min). Both groups show only sparse Iba1-positive cells, indicating no significant immune cell infiltration or inflammatory response (scale bar: 50 μm).

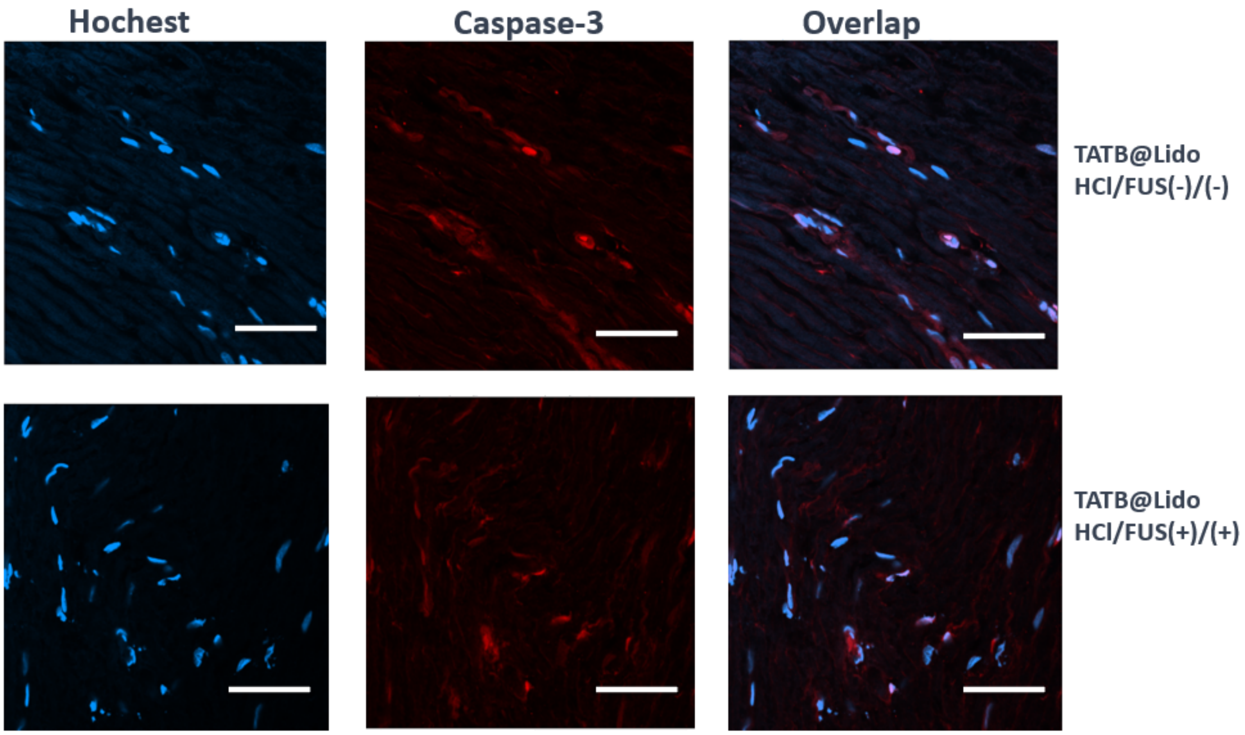


**Supplementary Fig. 10.** Representative immunofluorescence images of rat sciatic nerve sections 7 days after treatment, stained with Hoechst (blue, nuclei) and cleaved Caspase-3 (red, apoptotic marker). Top: TATB@Lidocaine HCl without FUS activation; Bottom: with FUS activation (1.5 MHz, 1.40 MPa, 8 min). Both groups exhibited minimal Caspase-3-positive staining, indicating no significant apoptosis in sciatic nerve tissue (scale bar: 50 μm).

### Supplementary Tables

**Supplementary Table 1.** Complete list of 246 compounds in the Focused Library used for machine-learning-based screening.

| Number | Compound name | Number | Compound name | Number | Compound name |
|---|---|---|---|---|---|
| 1 | Aceclofenac | 21 | Benzhydrocodone | 41 | Chlorobutanol |
| 2 | Acemetacin | 22 | Benzonatate | 42 | Chloroprocaine |
| 3 | Acetaminophen | 23 | Benzydamine | 43 | Chlorphenesin |
| 4 | Acetylsalicylic acid | 24 | Berotralstat | 44 | Chlorzoxazone |
| 5 | Alclofenac | 25 | Bromfenac | 45 | Choline magnesium trisalicylate |
| 6 | Alfentanil | 26 | Bronopol | 46 | Choline salicylate |
| 7 | Almotriptan | 27 | Bupivacaine | 47 | Cilostazol |
| 8 | Alverine | 28 | Buprenorphine | 48 | Cinchocaine |
| 9 | Alvimopan | 29 | Butalbital | 49 | Cisatracurium |
| 10 | Amitriptyline | 30 | Butamben | 50 | Clidinium |
| 11 | Amitriptyline | 31 | Butorphanol | 51 | Clobetasol propionate |
| 12 | Amphetamine | 32 | Butylscopolamine | 52 | Clomipramine |
| 13 | Amylocaine | 33 | Caffeine | 53 | Clonidine |
| 14 | Anileridine | 34 | Cannabidiol | 54 | Cocaine |
| 15 | Antrafenine | 35 | Capsaicin | 55 | Codeine |
| 16 | Articaine | 36 | Carisoprodol | 56 | Cyclobenzaprine |
| 17 | Atracurium besylate | 37 | Carprofen | 57 | Decamethonium |
| 18 | Baclofen | 38 | Celecoxib | 58 | Desflurane |
| 19 | Bazedoxifene | 39 | Chloral hydrate | 59 | Desipramine |
| 20 | Bendazac | 40 | Chlorcyclizine | 60 | Dexketoprofen |

| Number | Compound name | Number | Compound name | Number | Compound name |
|---|---|---|---|---|---|
| 61 | Dextropropoxyphene | 81 | Enflurane | 101 | Ganaxolone |
| 62 | Dezocine | 82 | Ephedrine | 102 | Glucosamine |
| 63 | Diamorphine | 83 | Epinephrine | 103 | Glycol salicylate |
| 64 | Diclofenac | 84 | Ethanol | 104 | Glycopyrronium |
| 65 | Dienogest | 85 | Etidocaine | 105 | Guaiacol |
| 66 | Diflunisal | 86 | Etodolac | 106 | Halothane |
| 67 | Difluprednate | 87 | Etomidate | 107 | Hexafluronium |
| 68 | Dihydrocodeine | 88 | Etoricoxib | 108 | Hexylcaine |
| 69 | Dimethyl sulfoxide | 89 | Eugenol | 109 | Hexylresorcinol |
| 70 | Disopyramide | 90 | Fenoprofen | 110 | Hyaluronic acid |
| 71 | Domperidone | 91 | Fentanyl | 111 | Hydrocodone |
| 72 | Doxacurium | 92 | Flavoxate | 112 | Hydromorphone |
| 73 | Doxepin | 93 | Flecainide | 113 | Hydroxyurea |
| 74 | Dronabinol | 94 | Flumazenil | 114 | Hyoscyamine |
| 75 | Droperidol | 95 | Flurbiprofen | 115 | Imipramine |
| 76 | Duloxetine | 96 | Fospropofol | 116 | Indecainide |
| 77 | Dyclonine | 97 | Gabapentin | 117 | Indomethacin |
| 78 | Edaravone | 98 | Gallamine triethiodide | 118 | Isoflurane |
| 79 | Elagolix | 99 | gamma-Hydroxybutyric acid | 119 | Isoprenaline |
| 80 | Eluxadoline | 100 | Gamolenic acid | 120 | Ketamine |
| Number | Compound name | Number | Compound name | Number | Compound name |
| 121 | Ketoprofen | 141 | Meloxicam | 161 | Morphine |
| 122 | Ketorolac | 142 | Menthyl salicylate | 162 | Nabilone |
| 123 | Lamotrigine | 143 | Meperidine | 163 | Nalbuphine |

| | | | | | |
|---|---|---|---|---|---|
| 124 | Levacetylmethadol | 144 | Mephentermine | 164 | Naloxegol |
| 125 | Levobupivacaine | 145 | Mepivacaine | 165 | Naloxone |
| 126 | Levomenthol | 146 | Metamizole | 166 | Nandrolone decanoate |
| 127 | Levorphanol | 147 | Metaraminol | 167 | Naproxen |
| 128 | Lidocaine | 148 | Metaxalone | 168 | Nepafenac |
| 129 | Linzagolix | 149 | Methadone | 169 | Nicoboxil |
| 130 | Lofexidine | 150 | Methocarbamol | 170 | Nimesulide |
| 131 | Lorazepam | 151 | Methohexital | 171 | Nitroglycerin |
| 132 | Lornoxicam | 152 | Methoxyflurane | 172 | Norethisterone |
| 133 | Loteprednol etabonate | 153 | Methyl nicotinate | 173 | Nortriptyline |
| 134 | Loxoprofen | 154 | Methyl salicylate | 174 | Oliceridine |
| 135 | Lubiprostone | 155 | Methyltestosterone | 175 | Orphenadrine |
| 136 | Meclizine | 156 | Metoclopramide | 176 | Oxaprozin |
| 137 | Meclofenamic acid | 157 | Metocurine iodide | 177 | Oxetacaine |
| 138 | Medifoxamine | 158 | Midazolam | 178 | Oxybuprocaine |
| 139 | Medroxyprogesterone acetate | 159 | Milnacipran | 179 | Oxycodone |
| 140 | Mefenamic acid | 160 | Mivacurium | 180 | Oxymetazoline |
| Number | Compound name | Number | Compound name | Number | Compound name |
| 181 | Oxymorphone | 201 | Pramocaine | 221 | Sufentanil |
| 182 | Ozanimod | 202 | Prasterone | 222 | Sugammadex |
| 183 | Pancuronium | 203 | Pregabalin | 223 | Sulindac |
| 184 | Parecoxib | 204 | Prilocaine | 224 | Tannic acid |
| 185 | Pentaerythritol tetranitrate | 205 | Procaine | 225 | Tapentadol |
| 186 | Pentazocine | 206 | Procaine benzylpenicillin | 226 | Technetium Tc-99m disofenin |
| 187 | Pentosan polysulfate | 207 | Promethazine | 227 | Tenoxicam |

| | | | | | |
|---|---|---|---|---|---|
| 188 | Pentoxifylline | 208 | Proparacaine | 228 | Thiamylal |
| 189 | Pentoxyverine | 209 | Propoxycaine | 229 | Thiethylperazine |
| 190 | Phenazopyridine | 210 | Pyrithione | 230 | Thiopental |
| 191 | Phenol | 211 | Remifentanil | 231 | Tiaprofenic acid |
| 192 | Phentolamine | 212 | Remimazolam | 232 | Tirofiban |
| 193 | Phenyl salicylate | 213 | Rocuronium | 233 | Tizanidine |
| 194 | Phenylephrine | 214 | Rofecoxib | 234 | Tolfenamic acid |
| 195 | Phenyltoloxamine | 215 | Ropivacaine | 235 | Tramadol |
| 196 | Pinaverium | 216 | Salicylamide | 236 | Triamcinolone |
| 197 | Pipecuronium | 217 | Samarium (153Sm) lexidronam | 237 | Trichloroethylene |
| 198 | Pipotiazine | 218 | Sevoflurane | 238 | Triflusal |
| 199 | Piritramide | 219 | Sildenafil | 239 | Trimebutine |
| 200 | Piroxicam | 220 | Succinylcholine | 240 | Trimethaphan |
| Number | Compound name | Number | Compound name | Number | Compound name |
| 241 | Trolamine salicylate | 243 | Vecuronium | 245 | Zavegepant |
| 242 | Tryptophan | 244 | Venlafaxine | 246 | Zucapsaicin |

**Supplementary Table 2.** The Training Dataset for machine learning models.

| Number | Name | loading_mol (%) | HOF | Encapsulation Method |
|---|---|---|---|---|
| 1 | Aspirin | 12.1 | HOF-TATB | DSM |
| 2 | Benzocaine | 26.7 | HOF-TATB | DSM |
| 3 | Benzocaine | 68.5 | HOF-101 | DSM |
| 4 | Benzocaine | 72.1 | HOF-102 | DSM |
| 5 | Carbamazepine | 28.6 | HOF-TATB | DSM |
| 6 | Carbamazepine | 71.3 | HOF-101 | DSM |
| 7 | Carbamazepine | 76.2 | HOF-102 | DSM |
| 8 | GM6001 | 31 | HOF-TATB | DSM |
| 9 | Ibuprofen | 38.3 | HOF-TATB | DSM |
| 10 | Ibuprofen | 30.1 | HOF-BTB | DSM |
| 11 | Ibuprofen | 64.5 | HOF-101 | DSM |
| 12 | Ibuprofen | 73.2 | HOF-102 | DSM |
| 13 | Isradipine | 30.8 | HOF-TATB | DSM |
| 14 | Isradipine | 16.7 | HOF-BTB | DSM |
| 15 | Isradipine | 53.2 | HOF-101 | DSM |
| 16 | Isradipine | 66.2 | HOF-102 | DSM |
| 17 | Tetracaine | 32.2 | HOF-TATB | DSM |

| | | | | |
|---|---|---|---|---|
| 18 | Tetracaine | 26.2 | HOF-BTB | DSM |
| 19 | Tetracaine | 65.9 | HOF-101 | DSM |
| 20 | Tetracaine | 76.5 | HOF-102 | DSM |
| 21 | DCZ | 12.8 | HOF-TATB | Diffusion |
| 22 | DHPG | 6.1 | HOF-TATB | Diffusion |
| 23 | Dopamine | 15 | HOF-TATB | Diffusion |
| 24 | Dopamine | 18 | HOF-101 | Diffusion |
| 25 | Dopamine | 43 | HOF-102 | Diffusion |
| 26 | L-dopa | 9.2 | HOF-TATB | Diffusion |
| 27 | L-dopa | 32.4 | HOF-101 | Diffusion |
| 28 | L-dopa | 41 | HOF-102 | Diffusion |
| 29 | MethylenebBlue | 8.6 | HOF-TATB | Diffusion |
| 30 | MethylenebBlue | 11.5 | HOF-BTB | Diffusion |
| 31 | Methylprednisolone | 3.1 | HOF-TATB | Diffusion |
| 32 | Methylprednisolone | 1.5 | HOF-BTB | Diffusion |
| 33 | Methylprednisolone | 41.4 | HOF-101 | Diffusion |
| 34 | Methylprednisolone | 60.3 | HOF-102 | Diffusion |
| 35 | RhodamineB | 14.2 | HOF-TATB | Diffusion |
| 36 | RhodamineB | 14.6 | HOF-BTB | Diffusion |

| | | | | |
|---|---|---|---|---|
| 37 | RhodamineB | 34.3 | HOF-101 | Diffusion |
| 38 | RhodamineB | 44.3 | HOF-102 | Diffusion |
| 39 | Scopolamine | 5.8 | HOF-TATB | Diffusion |
| 40 | Scopolamine | 24.8 | HOF-101 | Diffusion |
| 41 | Scopolamine | 36.6 | HOF-102 | Diffusion |
| 42 | Tetracaine | 16.7 | HOF-TATB | Diffusion |
| 43 | Tetracaine | 20.2 | HOF-BTB | Diffusion |
| 44 | Tetracaine | 42.9 | HOF-101 | Diffusion |
| 45 | Tetracaine | 45.2 | HOF-102 | Diffusion |

**Supplementary Table 3.** Antibodies used in this work.

| **Primary antibodies** | **Secondary antibodies** |
|---|---|
| Rabbit anti-Iba1<br>(1:500, 013-27691, Wako Chemicals) | Donkey anti-Rabbit, Alexa Fluor 594<br>(1:500, A32754, Invitrogen) |
| Rabbit anti-Cleaved Caspase-3<br>(1:500, 9661, Cell Signaling Tec.) | Donkey anti-Rabbit, Alexa Fluor 594<br>(1:500, A32754, Invitrogen) |
| | H&E staining kit<br>(ab245880, Abcam) |
| | Hoechst 33342<br>(1:5000, 17535, AAT Bioquest ) |
